# Tumor-on-Chip as a Personalised Platform for Rapid Drug-Testing in Breast Cancer

**DOI:** 10.1101/2025.10.26.684605

**Authors:** Clara Helal, Léa Pinon, Juliette Jin, Niccolo Schintu, Ana Vazquez, Martin Nurmik, Gianni Antonelli, Maxime Simon, Elodie Montaudon, Laura Sourd, Heloise Derrien, Gérard Zalcman, Angela Bellini, Andreia Goncalves, Anne Vincent-Salomon, Toulsie Ramtohul, Elisabetta Marangoni, Maria Carla Parrini, Luc Cabel, Stéphanie Descroix

## Abstract

Breast cancer (BC) remains one of the most common malignancies worldwide and continues to pose major therapeutic challenges, emphasizing the need for functional models that can inform fast treatment selection. Currently, patient-derived xenografts (PDXs) and organoids (PDOs) are valuable biological models for functional precision oncology; however, variable success rates of establishment and prolonged timelines limit their clinical application for real-time drug testing. To overcome this, we developed a Tumor-on-Chip (ToC) platform that enables functional drug sensitivity profiling within a clinically actionable 4-day timeframe. We compared ToC models with PDX results, and demonstrated high reproducibility and strong concordance with *in vivo* PDX responses. Drug sensitivity was correctly identified *ex vivo* in 88% of PDX-responsive cases, while resistance was detected in 91%, with no false positives at clinically relevant drug concentrations. To facilitate clinical translation, we engineered a custom microfluidic chip optimized for minimal breast cancer biopsy material, yielding results similar to those obtained from resection samples. We ultimately demonstrated the proof-of-concept for applying this platform to patient samples as a further tool for guiding clinical decision-making and discriminated between resistance and sensitive profiles among patients. These findings demonstrate the feasibility and translational potential of ToC models for personalized drug profiling in breast cancer, laying the groundwork for their integration into real-time clinical decision-making workflows.

## Introduction

Breast cancer (BC) is the most prevalent malignancy and remains the second leading cause of cancer-related mortality among women worldwide (1). It encompasses a biologically heterogeneous disease, broadly categorized into three main molecular subtypes, each associated with distinct prognostic implications and therapeutic vulnerabilities: hormone receptor-positive/HER2-negative (HR+/HER2–, ∼ 70%), HER2-positive (∼ 15%), and triple-negative breast cancer (TNBC, ∼ 15%) (2, 3). While patients with primary tumor HR+/HER2– BC are treated with endocrine-based therapies (4); and HER2+ BC are treated with anti-HER2 therapies (5), chemotherapy remains the mainstay of treatment in the metastatic setting of any BC subtype, either after disease progression or in combination with targeted therapies (6, 7). Standard chemotherapeutic agents used in BC management non-exhaustively include taxanes, anthracyclines, carboplatin, and capecitabine (8). However, the optimal selection of cytotoxic agents remains undefined, as response rates are highly variable, ranging from 10 to 50% – depending on the agent, clinical context, and treatment setting – and no robust predictive biomarker is currently available to guide chemotherapy selection. As a result, therapeutic decisions are largely empirical, exposing patients to potentially ineffective regimens and avoidable toxicities. In recent years, valuable advances in sequencing technologies have enabled comprehensive profiling of molecular signatures – including genomic, transcriptomic, epigenomic, and proteomic data – fueling a strong push to translate precision medicine into clinical oncology practice (9–13). Genomics-driven approaches have deepened our understanding of pathways underlying tumorigenesis and cancer progression, facilitating the identification of oncogenic driver mutations (14). Consequently, considerable efforts have been directed towards individualizing treatment by matching therapies to patients’ molecular alterations (15). However, most of the identified genomic alterations are not therapeutically actionable and lack consistent predictive power for guiding treatment decisions at the individual level, highlighting ongoing challenges of current molecularly-guided strategies (16). One promising alternative relies on patient-specific tumor models, so-called *avatars*, which aim to recapitulate tumor biology and individual patient-specific factors (17, 18). By directly measuring drug responses in patient-derived tumor cells, functional precision medicine circumvents the limitations of genomics alone, enabling the discovery of new treatment options and repurposing opportunities in realtime (19).

Patient-derived xenograft (PDX) models have long been considered among the most robust *in vivo* avatar models for cancer research. By preserving the histological and genomic landscape of the patient’s tumor (17, 20, 21), PDXs have enabled major advances in both fundamental and translational oncology. Several studies have further demonstrated a strong concordance between PDX responses and clinical outcomes (22–25), establishing them as the historical gold-standard surrogate for predicting patient response to cytotoxic drugs, including in breast cancer. However, the variable engraftment efficiency, high costs, and most critically, the extended time required to obtain drug sensitivity profiles (often several months) preclude their use in routine clinical practice (17). Thus, innovative technologies that can rapidly generate patient-specific, high-throughput–compatible models are needed to provide clinically actionable insights within a relevant therapeutic timeframe.

One such valuable approach is the use of patient-derived organoids (PDO), which provide high-throughput and scalable *in vitro* platforms. Numerous studies have successfully generated PDOs from a wide range of human malignancies, including breast, colorectal, renal, ovarian, pancreatic, and liver cancers (26). These PDO models enable the capture of cellular heterogeneity and spatial architecture of tumors (27), providing valuable platforms for investigating tumor initiation and disease progression (28). PDOs have also demonstrated strong predictive value for drug response across various cancer types (29), supporting their integration as translational tools for therapy selection within a functional precision medicine framework (30). However, two major limitations currently hinder their translation into routine clinical practice. First, the efficiency of PDO establishment is inconsistent and often suboptimal, with reported success rates ranging from 17 to 90% depending on tumor subtype, disease aggressivity, and sample cellularity (31–33). Second, the extended timeframe required to establish and expand PDO cultures (several weeks), including in BC (12), strongly limits their applicability for real-time drug screening (34).

To address these limitations, Tumor-on-Chip (ToC) technology has emerged as a compelling alternative that may ultimately complement traditional preclinical models (35). These miniaturized 3D platforms are engineered to replicate *ex vivo* the essential features of living tissues, including the cellular diversity of the tumor microenvironment (TME) and the physico-chemical parameters of the extracellular matrix (ECM) (36, 37). Nevertheless, drug-testing using ToC technology remains in its early stages, with only a few groups exploring the feasibility of using patient-derived tumor cells in microfluidic systems to predict therapeutic responses (38–40). A few techniques have emerged towards cuboids, *i*.*e*., microdissection from intact tissues of biopsies, and tumor slice culture, to provide a high-throughput microfluidic platform for drug screening (41, 42). In the context of validating the correlation between *ex vivo* techniques and *in vivo* outcomes (43), a multiplexed ToC model of colorectal cancer derived from PDX was successfully established within only four days, demonstrating on three models a rather good *in vitro*–*in vivo* correlation with corresponding PDX responses (44). Although very encouraging, the few existing studies investigating the concordance between ToC responses and patient outcomes – primarily in lung, colorectal, and pancreatic cancer (39, 45) – remain proof-of-concept studies conducted in small patient cohorts, and are not powered to establish statistically robust correlations. To date, no breast cancer–specific ToC model has been clinically evaluated for its predictive accuracy.

In this study, we explore the potential of breast cancer ToC platform designed for *ex vivo* prediction of therapeutic efficacy within a clinically actionable timeframe, with the long-term objective of supporting real-time treatment decisions. To this end, using PDX as patient source, (i) we validated our breast cancer ToC model against matched PDX models – the current, albeit imperfect, gold standard for patient outcomes prediction – to assess drug response sensitivity and (ii) evaluated the practical feasibility of generating patient-specific ToC directly from fresh tumor resections and core needle biopsies. We established PDX-derived ToC across six breast cancer PDX models and characterized their drug sensitivity and resistance profiles using both chemotherapies and antibody-drug conjugates (ADCs), demonstrating the ability of the ToC to capture the patient-specific drug response. We then compared the drug sensitivity profile between *ex vivo* ToC and their corresponding *in vivo* PDX, revealing strong concordance. Building on this, we leveraged the microfluidic technology to develop and implement a new microfluidic chip designed specifically for patient samples, with a focus on low-input biopsy samples, to assess patient-specific drug sensitivity. Our findings lay the groundwork for the clinical translation of patient-derived ToC platforms as functional precision oncology tools to guide individualized treatment strategies in metastatic breast cancer.

## Results

### PDX-derived ToC technology enables the assessment of drug responses on patient-derived cells in BC

To capture the heterogeneity of BC patients, we focused on the three main clinical subtypes: triple negative (ER-, PR-, HER2-), hormone receptor-positive (ER+ and/or PR+), and HER2-positive (**Fig. 1A**). Accordingly, we selected a panel of six PDX models of BC established from either primary tumors or metastatic lesions (**Fig. S1**). Contrary to recent ToC techniques (42), we isolated individual tumor cells from PDXs to develop PDX-corresponding ToC models. Following PDX excision, tumor cells were exclusively isolated within half a day by lysing red blood cells, removing debris using a density gradient, and removing dead and murine cells using magnetic cell sorting (MACS) (**Fig. 1B**). The purified tumor cells were then embedded in a collagen I matrix and introduced into the central channel of a microfluidic chip (**Fig. 1C**) with a success rate >99% (n=86 ToC models established out of 88 PDX tumors). To assess drug effects across the different BC models, cell viability was monitored longitudinally using a live/dead assay under varying drug classes and concentrations added to the lateral chambers of the chip after gel polymerization (**Fig. 1D**).

**Fig. 1.**
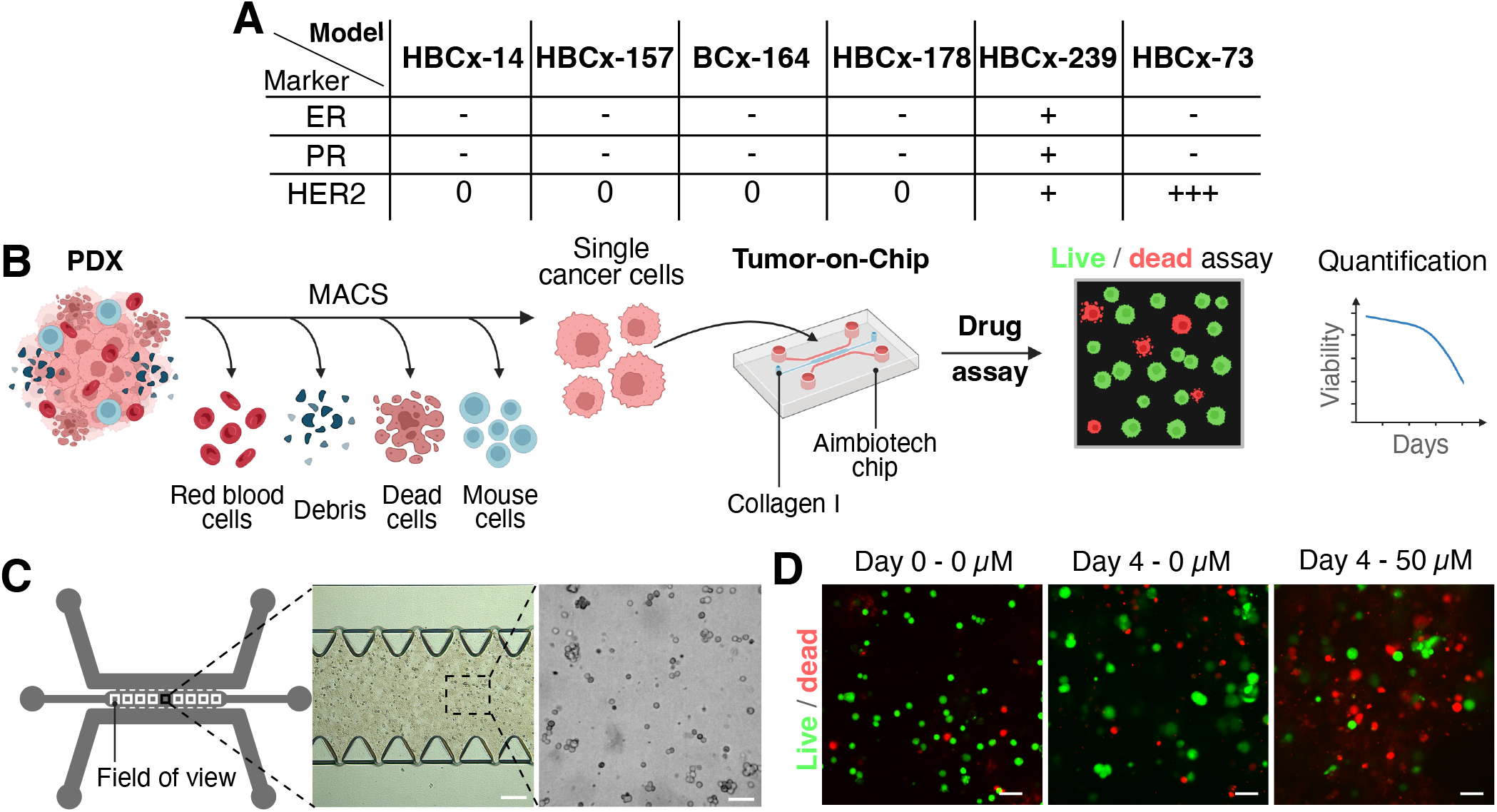
Strategy for assessment of drug responses using PDX-derived Tumor-on-chip. (A) Table recapitulating the characteristics of the PDX models used in the study, including the IHC expression of breast cancer markers. ER = Estrogen Receptor, PR = Progesterone Receptor, HER2 = Human Epidermal Growth Factor Receptor 2. (B) Schematic of the experimental pipeline. Tumor cells isolated from PDXs by magnetic cell sorting (MACS) were embedded in collagen I gel and seeded into the microfluidic chips. Cell viability was monitored over time using a live/dead assay and quantified. (C) Schematic of the chip layout. For each condition, nine fields of view were imaged along the central microfluidic channel. Representative brightfield images of breast cancer cells embedded in collagen gel at low (4X) and higher magnification (10X). Scale bars: 200 µm (left) and 50 µm (right). (D) Confocal images of live/dead assays in HBCx-14 cancer cells before and 4 days after cisplatin exposure. Scale bar: 30 µm.

The ToC platform was first optimized using the HBCx-14 model by testing a range of collagen properties, including ECM stiffness and cell crowding, given that the TME is typically stiffer (46) and denser in cells than healthy tissues (47). Overall, tumor cell viability decreased in a dose-dependent manner upon cisplatin treatment, regardless of collagen I concentration (ranging from 2.5 mg/mL to 8 mg/mL) (**Fig. S2A**) or cell embedding density (10^3^ to 20.10^3^ cells/µL) (**Fig. S2B**). No statistically significant association was observed between these parameters and drug response to cisplatin. Based on these findings and supported by previous literature, we selected a collagen concentration of 2.5 mg/mL and a cell density of 4×10^3^ cells/µL as the standard condition for all subsequent experiments (48–51). Next, to select the most suitable imaging strategy, endpoint analysis (different chips imaged at successive timepoints) or live imaging (continuous monitoring of the same chip) were compared over four days using the HBCx-14 ToC model and three drugs including cisplatin, paclitaxel and 5FU (**Fig. S2C** and **S2D**). Despite some differences due to the varying sensitivities of the methods used to quantify viability (endpoint live/dead assay or caspase live reporter), treated cells exhibited similar viability trends in endpoint analysis compared to live imaging, both in terms of dose-response and time-course. Therefore, the endpoint strategy was selected for subsequent experiments as it facilitates the parallel testing of multiple conditions, maximizing throughput and offering greater ease of use.

Overall, the live/dead assay applied to PDX-derived ToC models provides a reliable assessment of breast cancer cell viability under drug treatments within a timeframe of less than one week.

### ToC enables rapid *ex vivo* multiple drug screening in patient-derived tumor cells

To investigate the effects of treatment in BC, PDX-derived tumor cells of the HBCx-14 were exposed in ToC to agents with distinct mechanisms of action: 5FU (nucleic acid synthesis inhibition); carboplatin and cisplatin (DNA cross-linking); and paclitaxel (microtubule stabilization and mitotic arrest), each ultimately resulting in cell death. After extracting and isolating tumor cells, we monitored the response of the HBCx-14 model to these agents in the ToC assay across a harmonized concentration range from 1 to 50 µM. This approach allowed us to determine the minimal exposure duration required to detect differential drug sensitivity *ex vivo*, thereby facilitating direct comparison with the corresponding *in vivo* treatment timeframes. A significant dose-dependent reduction in cell viability was observed by day 4 following treatment with cisplatin, carboplatin, and paclitaxel, while 5FU showed no effect at any concentration (**Fig. 2A**), showing that a 4-day window is sufficient to discriminate sensitivity profiles in PDX-derived ToC. Based on the 4-day drug assay values, we generated a heatmap of relative cell viability for this model, providing a concise overview of drug sensitivity across all conditions (**Fig. 2B**). We performed Alamar Blue assay as a complementary metabolic endpoint cross-validation of the live-dead assay, and similar patterns of drug sensitivity were obtained, excluding any assay-specific bias in ToC (**Fig. S3**).

**Fig. 2.**
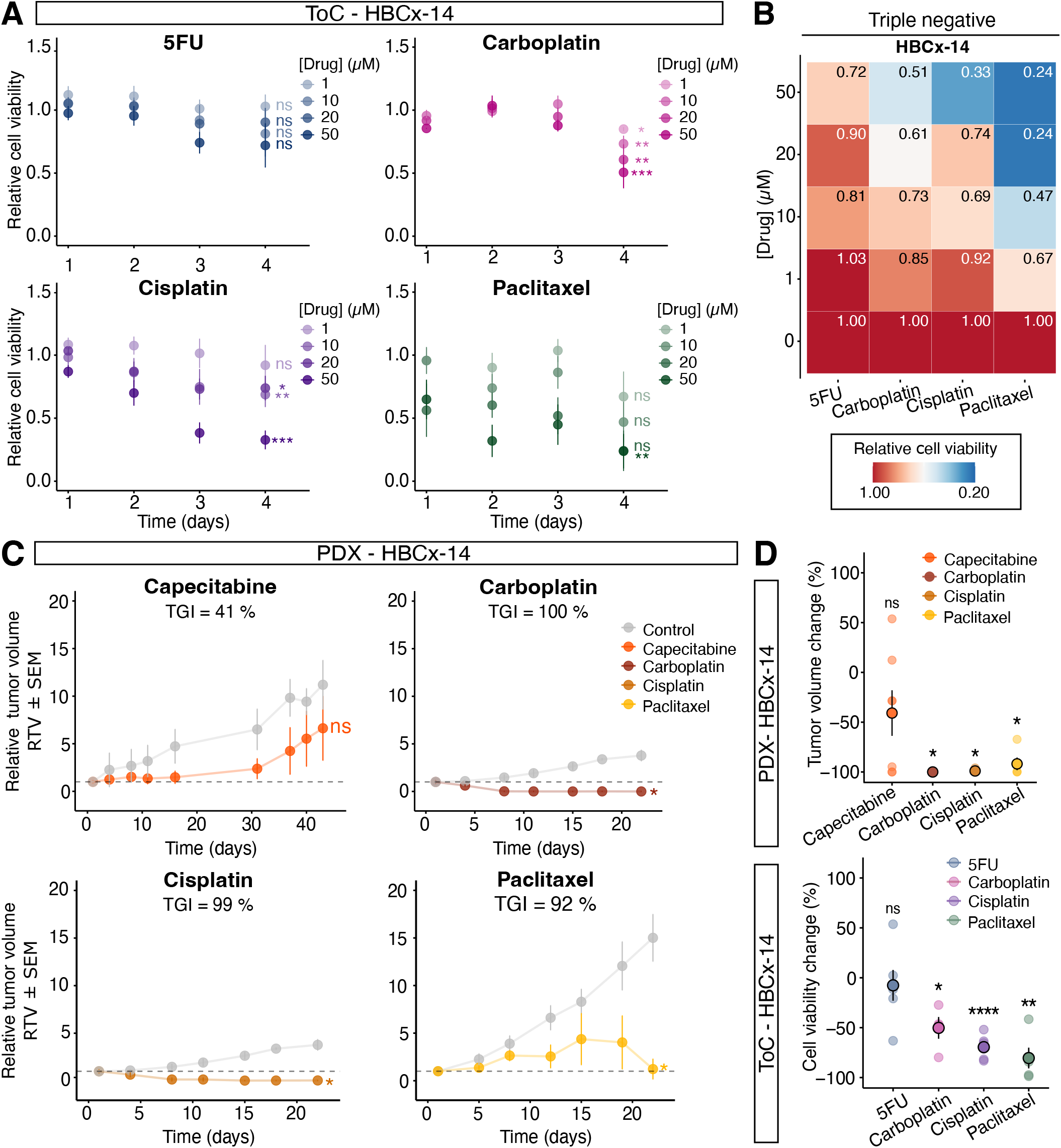
ToC enables rapid *ex vivo* multiple drug screening in patient-derived tumor cells and recapitulates *in vivo* drug responses of HBCx-14 PDX models. (A) Graphs representing the kinetics of cell viability over time for the HBCx-14 model treated with 5FU, cisplatin, carboplatin, and paclitaxel from 1 to 50 µM. In all conditions, individual data were normalized by the initial point (Day 0), and with the control condition (concentration = 0) at each day. Each point represents the mean of at least 3 independent experiments, and the error bar corresponds to a SD. All statistics were performed using the Wilcoxon test. ns = p-value > 0.05, * = p-value *≤* 0.05, ** = p-value *≤* 0.01, *** = p-value *≤* 0.001, **** = p-value *≤* 0.0001. (B) Heat maps of relative cell viability at day 4 for the HBCx-14 tumor model. Colors indicate the drug response. Sensitivity and resistance are indicated in blue and red, respectively. (C) Kinetics of PDX tumor growth *in vivo* in the HBCx-14 model. Tumor volumes (mm^3^) were measured longitudinally; each point represents the mean of at least three independent mice, and error bars indicate the SEM. Treated groups are shown as colored curves and untreated controls as a gray curve. Treatment regimens were: capecitabine, 540 mg/kg, five times per week per os; carboplatin, 90 mg/kg, once every three weeks IP; cisplatin, 6 mg/kg, once every three weeks IP; and paclitaxel, 25 mg/kg, once per week IP. TGI = tumor growth inhibition = [1-(RTV_*treated*_/RTV_*control*_)]*100 at day_*max*_. For intermediate TGI (60–90%), histogram shows the relative mean tumor volume change (%VC) of the treated group, illustrating resistance to treatment. (D) Comparison of tumor volume change (VC, %) in PDX and cell viability change (%) in the corresponding ToCs in the HBCx-14 model. For Tumor-on-Chip, cell viability change values correspond to treatment with 50 µM of each drug.

We then evaluated the predictive performance of ToC against PDX, the current gold standard surrogate for patient response. To this end, tumor growth was concurrently monitored over time in mice (**Fig. S4A-B**). Relative tumor growths of HBCx-14 tumor-grafted mice, treated and non-treated with different drugs, are displayed in **Figure 2C**. Platinum-based drugs and paclitaxel induced a significant reduction of tumor volume compared to control conditions, indicating an *in vivo* sensitivity to carboplatin, cisplatin, and paclitaxel, whereas capecitabine (the oral prodrug of 5FU) had no significant effect after 40 days of exposure, similarly to those obtained in ToC. Such observations are confirmed by the similar tendency of the percentage change in tumor growth (*in vivo*) and cell viability (on-chip, *ex vivo*) relative to untreated controls (**Fig. 2D**), suggesting that ToC correctly mirrors the matched PDX model, while reducing response time to 4 days.

Next, to further discriminate sensitivity and resistance profiles in ToC, we determined a quantitative threshold of cell viability based on the responses of the HBCx-14 model in PDX. To this end, PDX responses were classified following two standard metrics of preclinical modeling of antitumor activity *in vivo* (52): the tumor growth inhibition (TGI) and the percentage of tumor volume change (%VC) (see Materials & Methods section). PDX models were classified resistant to treatment when TGI ≤ 60% or intermediate (60-90%) with progressive disease (%VC *>* +35%); and sensitive to treatment when TGI ≥ 90% or intermediate (60-90%) without progressive disease (%VC ≤ +35%). Consistent with the response patterns observed in the matched HBCx-14 model *in vivo*, we considered a reduction greater than 33% in relative tumor cell viability as a biologically relevant threshold to discriminate drug sensitivity profiles in the ToC assay.

### *Ex vivo* ToC assays recapitulate *in vivo* drug responses of breast cancer PDX models

To further use the ToCs as a predictive platform, we applied the ToC assays across the wider panel of different PDX models (**Fig. 1A**) using the predefined threshold. To preserve biological relevance, only standard-of-care agents routinely employed in the clinical management of breast cancer were tested (53): 5FU (the active metabolite of capecitabine), carboplatin, and paclitaxel. As expected from differences in tumor types and drug mechanisms, the ToC assay revealed heterogeneous responses across models after 4 days of culture (**Fig. 3A** and **S5**). Specifically, HBCx-164 and HBCx-157 exhibited a dose-dependent sensitivity to both carboplatin and paclitaxel, whereas the triple-negative model, HBCx-178, was resistant to these agents across all tested doses. The HR+/HER2+ model (HBCx-239) was resistant to all treatments, and the HER2 3+ model (HBCx-73) demonstrated selective sensitivity to 5FU. Altogether, these findings highlight the ToC platform’s capacity to capture subtype-specific drug sensitivities and resistances, demonstrating robust discrimination of treatment responses across diverse BC models within only 4 days.

**Fig. 3.**
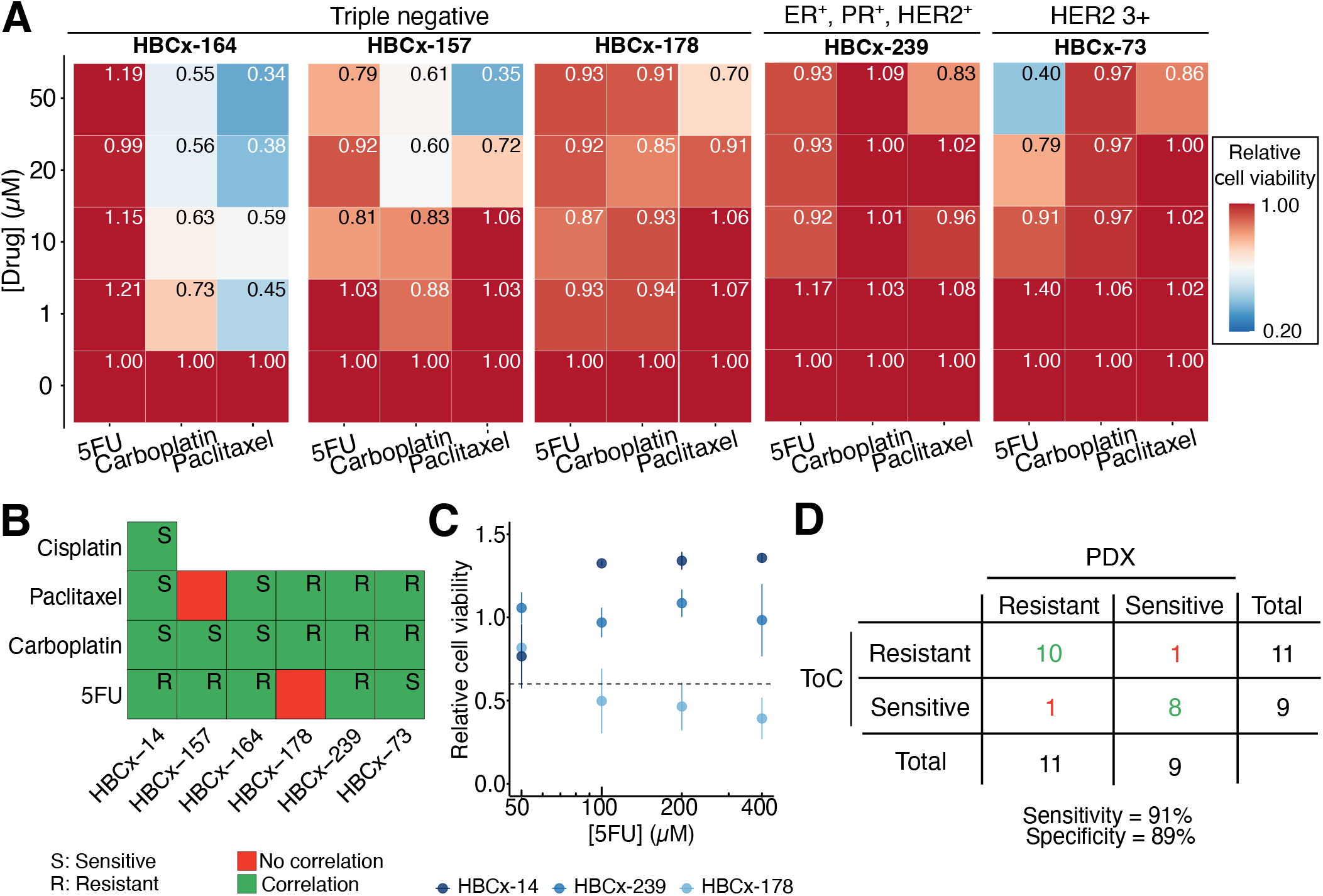
Drug sensitivity profiles identified *ex vivo* in ToC are concordant with *in vivo* responses. (A) Heat maps of relative cell viability at day 4 for the five other PDX models of the panel, treated with 5-FU, carboplatin and paclitaxel. Colors indicate the drug response. Sensitivity and resistance are indicated in blue and red, respectively. (B) Correlation between tumor volume change (PDX) and cell viability change (Tumor-on-Chip) across the panel of PDX models of BC. (C) Concordance of drug responses across each model tested. PDX models were classified resistant to treatment *in vivo* when TGI ≤ 60% or intermediate (60-90%) with progressive disease (%VC *>* +35%); and sensitive to treatment when TGI ≥ 90% or intermediate without progressive disease (%VC ≤ +35%). PDX-derived ToC models were considered sensitive to treatment *ex vivo* if a reduction of at least 33% in normalized tumor cell viability compared to baseline was observed at day 4. (D) Relative cell viability of HBCx-14, HBC-178, and HBCx-239 treated with concentrations of 5FU ranging from 50 to 400 µM.(E) Contingency table evaluating the diagnostic performance of the ToC assay. 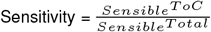and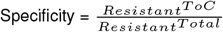.

To evaluate the predictive performance of the *ex vivo* ToC assay, we compared its readouts with the corresponding *in vivo* responses of the matched PDX models. Complete tumor growth kinetics for all conditions are provided in **S6A-C**. *In vivo* tumor growth and *ex vivo* cell viability were largely concordant across models, consistently identifying HBCx-14, HBCx-164, and HBCx-157 as sensitive to carboplatin; HBCx-14 and HBCx-164 to paclitaxel; HBCx-178 and HBCx-239 as resistant to these agents; and HBCx-73 as sensitive to 5FU (**Fig. 3B** and **Fig. S7**). Across all models and treatments, two discrepancies were observed. First HBCx-157, which is a model resistant to paclitaxel *in vivo*, was classified as sensitive in the ToC assay at 50 µM. Second, HBCx-178, a model known to be sensitive to capecitabine *in vivo*, was categorized as resistant to 5FU in the ToC assay at 50 µM (**Fig. S7**). Notably, HBCx-157 treated with paclitaxel at 1 and 10 µM – within the range of patient plasma levels (54) – remained resistant, suggesting that this model only responds only at supra-clinical concentrations, unlike the other models that remain resistant even at these high doses. To investigate the 5FU discrepancy, ToC assay was repeated at concentrations approaching the reported maximal plasma levels of 5FU in patients (54) (up to 400 µM). From a 5FU concentration of 100 µM, HBCx-178 restored sensitivity (**Fig. 3C**). In contrast, models known to be resistant to 5FU *in vivo*, including HBCx-14 and HBCx-239, remained refractory even at the highest concentration tested, further validating the assay efficiency.

Overall, when a drug demonstrated efficacy *in vivo*, the ToC assay (using 50 µM on day 4) correctly identified sensitivity *ex vivo* in 88% of cases (7 out of 8 drug–PDX combinations, across 6 PDX models treated with 3 clinically relevant agents each, plus cisplatin in HBCx-14). Conversely, drug resistance was consistently detected in 91% of cases (10 cases out of 11), highlighting the reliability of the ToC assay for distinguishing responders from non-responders (**Fig. 3D**).

Altogether, these results demonstrate that tumor-on-chip models closely match drug responses observed in the PDX models, achieving both high sensitivity and specificity, while delivering results with at least a 5-to 10-fold faster turnaround time compared to *in vivo* models.

### ToC models allow mechanistic study of complex emerging ADC anticancer therapies

Antibody–drug conjugates (ADCs) represent a paradigm shift in BC therapeutic management, combining selectivity with potent cytotoxic payloads (55). Building on the strong *ex vivo* – *in vivo* concordance observed with chemotherapy agents, we thus extended our evaluation to ADCs. We tested trastuzumab–deruxtecan (T-DXd) in two HER2-positive models: HBCx-239 (HER2 1+) and HBCx-73 (HER2 3+); and one HER2-negative model known to be resistant to T-DXd *in vivo*, the HBCx-264 (**Fig. 4A**).

**Fig. 4.**
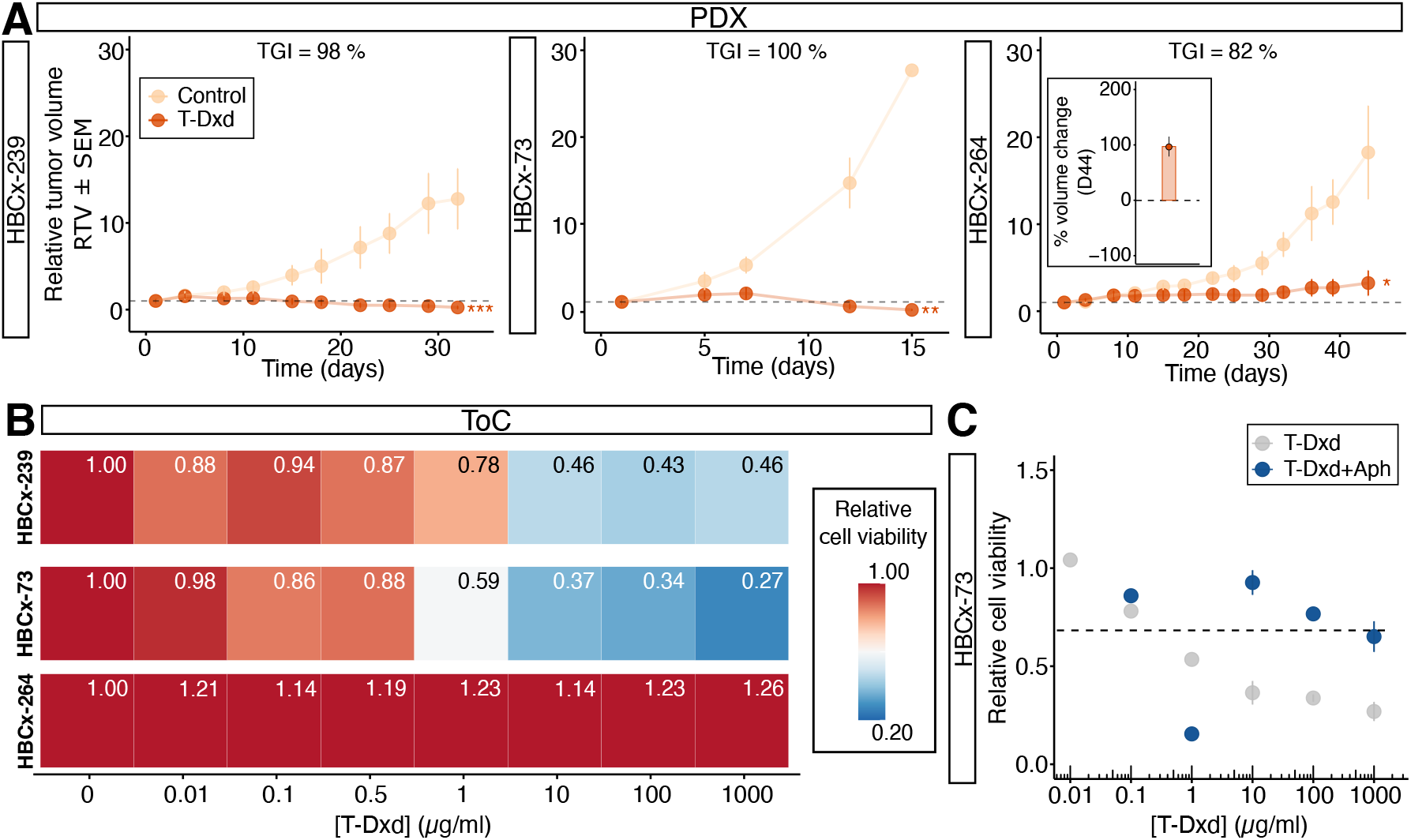
Tumor-on-chip models allow the mechanistic study of ADC strategy. (A) Graphs representing the kinetics of tumor volume growth in PDX for the HBCx-239, HBCx-73, and HBCx-264 BC models treated with T-DXd at 10 mg/kg. Each point represents the mean of at least 3 independent experiments. (B) Heat maps of relative cell viability at day 4 for HBCx-239, HBCx-73, and HBCx-264 BC models. Red indicates higher resistance, and blue shows a greater sensitivity to treatment. (C) Relative cell viability of HBCx-73 tumor models treated with T-DXd ± aphidicolin, an inhibitor of polymerase DNA to stop cell replication at the G1-phase.

Sensitive profiles were quantified on-chip within 4 days, once again underscoring the rapid readout capability of the ToC assay compared with the 15–45 days typically required for conventional *in vivo* mouse efficacy studies. Tumor cells were exposed to T-DXd concentrations ranging from 0.01 µg/mL to 1000 µg/mL, spanning the minimal payload (Dxd) release to 10-fold the maximal intact drug (T-DXd) concentration observed *in vivo*, respectively (56, 57). Both HER2-positive tumor models exhibited a dose-dependent decrease in viability (**Fig. 4B**), while HBCx-264 remained unresponsive, demonstrating the ToC platform’s ability to also faithfully reflect *in vivo* sensitivity and resistance profiles under ADC treatment. Notably, HBCx-73 displayed enhanced sensitivity, consistent with its higher HER2 expression (58). Furthermore, the sensitivity of HBCx-239 and HBCx-73 is marked at 10 µg/mL (**Fig. 4C**), corresponding to the T-DXd concentration at the end of a dosing cycle in patients (57), suggesting that the efficacy dose observed in ToC reflects the clinically relevant drug exposure.

Leveraging the versatility of the ToC platform, we investigated one of the facets of T-DXd’s mechanism of action. We hypothesized that limited proliferation of cancer cells on-chip could reduce sensitivity to T-DXd. T-DXd combines a monoclonal antibody targeting HER2 with a topoisomerase I inhibitor, which requires active DNA replication for optimal efficacy. To test this, we pretreated cells with aphidicolin, a DNA replication inhibitor which blocks the cell cycle in S phase and observed a reduced T-DXd-cytotoxicity in the HBCx-73 model in the ToC assay (**Fig. 4C**). Notably, sensitivity was restored at supra-clinical concentrations (57), suggesting alternative replication-independent mechanisms, such as transcriptional interference.

Altogether, these findings highlight the potential of ToC to functionally stratify patients for ADC therapies within a 4-day actionable timeframe, and also reveal mechanistic determinants of drug sensitivity, including receptor expression and proliferation status.

### Patient-derived ToC models can be generated using minimal cell input from biopsy samples

While PDX excisions yield abundant tumor material (on average 4×10^6^ cells per tumor of approximately 1500 mm^3^), they only partially reflect the clinical situation. In clinical practice, patient biopsies – obtained through minimally invasive procedures before treatment selection – are typically used to guide therapeutic decisions. Therefore, to evaluate the feasibility of our model in a clinically realistic setting, we first generated ToC models from freshly simulated biopsies performed *ex vivo* immediately upon arrival of PDX tumor samples in the laboratory (**Fig. 5A**). PDX tumors were biopsied *ex vivo* 3 or 4 times depending on the tumor size, using an 18G (*i*.*e*., D=1.02 mm) core-needle as routinely employed in clinical practice. This sampling strategy was designed to ensure adequate cellular yield for downstream analyses while respecting clinical routine constraints, allowing us to isolate 100×10^3^ to 720×10^3^ tumor cells from PDX tumors (**Fig. S8A**), compared to the 1 to 5 million cancer cells obtained from the whole PDX.

**Fig. 5.**
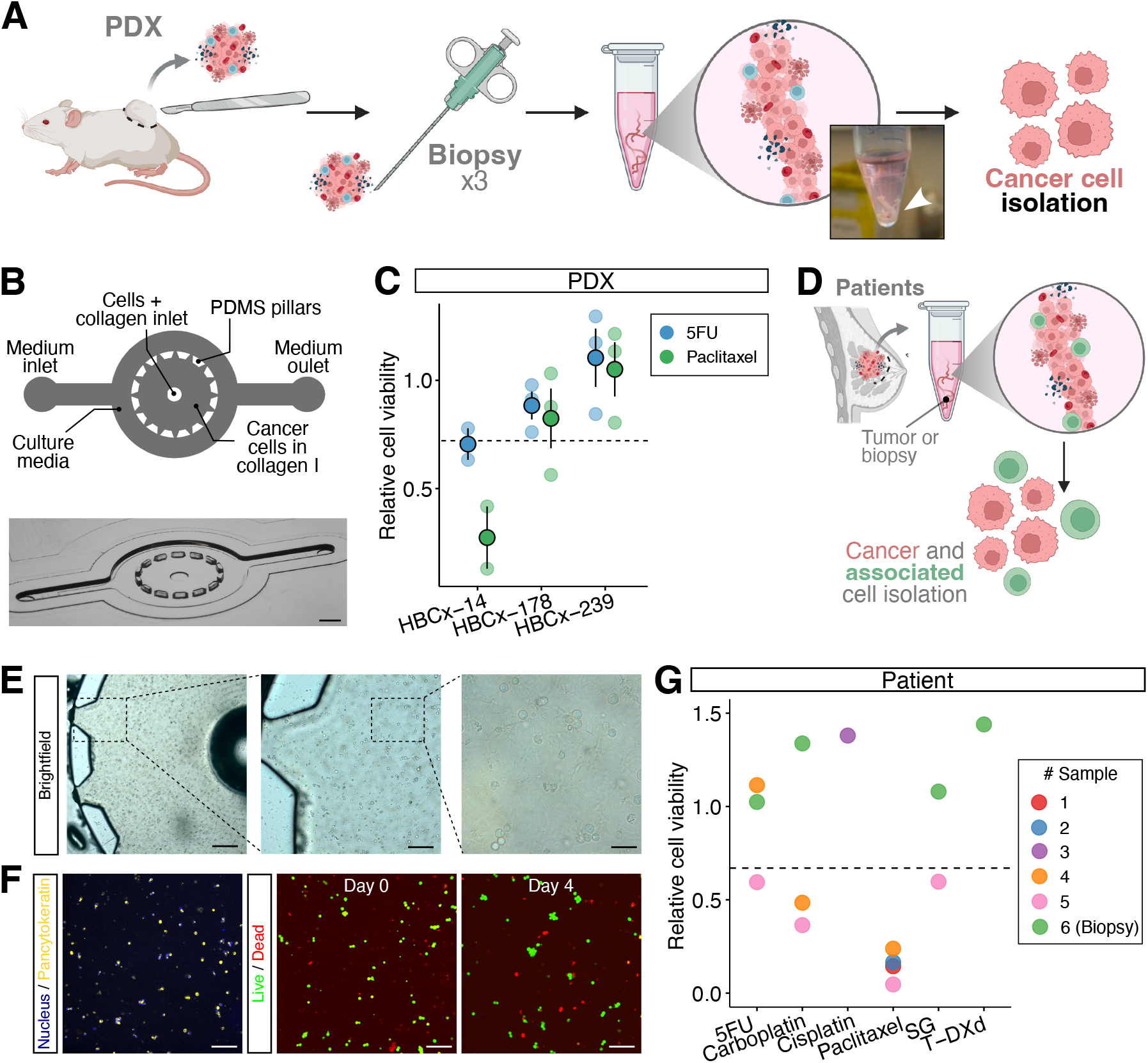
Patient-derived ToC models are established using minimal cell input from biopsy samples. (A) Schematics of the experimental pipeline of biopsies in PDX. Biopsies were performed *ex vivo* on surgical specimens of PDX. Tumor cells are isolated by removing murine cells (from PDX), red blood cells, and debris. (B) Representative images of a PDX tissue. (C) Representative image and schematics of the custom-chip for biopsies (top). Mold of the custom-made devices (bottom). (D) Cell viability of PDX biopsies in custom chips, treated with 5FU and paclitaxel at 50 µM. Values normalized to Day 0 and untreated condition at Day 4; dashed line = 33% efficacy threshold. (E) Schematics of the extraction of patient BC tissue and cell isolation. (F) Images of PDMS chips seeded with patient cells at different magnifications. Scale bars: 100, 50, 20 µm from left to right. (G) Confocal staining of patient-derived BC; nuclei and pancytokeratin (left) and live/dead assay (right) of sample #3 to paclitaxel (1 µM). (H) Relative cell viability of patient-derived cells after 4-day treatment with 5FU, carboplatin, cisplatin, paclitaxel (50 µM), sacituzumab-govitecan (SG, 1000 µM), and trastuzumab-deruxtecan (T-DXd, 1 µM).

Because biopsy-derived samples yield significantly fewer tumor cells than whole-tumor excisions, optimizing cell usage within the assay became a key priority to preserve analytical robustness despite limited input. To this end, we designed a custom PDMS-based microfluidic device featuring a 2.5 µL circular central chamber with a single inlet for seeding tumor cells embedded in collagen I, ensuring zero dead volume and accommodating approximately 10^4^ tumor cells per chip. This chamber is surrounded by a concentric medium channel with one inlet and one outlet for cell culture medium and drug delivery (**Fig. 5B**). Compared to commercial chips, our device requires 4-fold less matrix volume to fill the central channel, thereby reducing the initial number of cancer cells needed by 4-fold while maintaining the same cell concentration.

To validate the suitability of our custom chips for drug response assays, we performed the same cell viability assay on the biopsies-derived ToC from PDX tumors treated with 5FU and paclitaxel at 50 µM (**Fig. 5C**). The biopsy-derived HBCx-14 model was resistant to 5FU and sensitive to paclitaxel, consistent with the previous drug sensitivity profiles previously obtained from larger PDX samples and commercial chips (**Fig. 2B** and **Fig. 3A**). Similarly, biopsy-derived ToC from HBCx-239 and HBCx-178 exhibited resistance to both agents, while HBCx-73 showed selective sensitivity to 5FU. These results closely match the ToC sensitivity profiles obtained under conventional conditions, confirming that our custom biopsy-adapted ToC platform faithfully mirrors PDX drug-response profiles using a maximum of 10^4^ cells/chip - within a clinically relevant 4-day timeframe. This positions our platform as a powerful tool for evaluating prospective patient-specific treatments.

We next applied this approach to a primary human tumor samples to extend the clinical applicability of our assay (**Fig. 5D**). Surgical specimens (samples #1 to #5) of BC deemed unnecessary for diagnosis were retrieved and processed immediately to establish patient-derived ToC models (**Fig. 5E**). Tumor cells were seeded into the biopsy-chip at 10^4^ cells per chip and exposed to treatments at different concentrations. To confirm tumor cell identity on-chip, chips were fixed and subsequently stained with pancytokeratin, and a live/dead assay was performed (**Fig. 5F**). Cell viability relative to untreated controls was assessed after 4 days using a live/dead assay, revealing a significant decrease in cell viability. These results suggest that the tumors #1 and #2 were sensitive to paclitaxel and that the corresponding chemonaive patients may be responsive to this treatment (**Fig. 5G**). The tumor #3 is highly resistant to cisplatin. The chemonaive tumor #4 responds to carboplatin and paclitaxel, but is resistant to 5FU, while the chemonaive tumor #5 seems to respond to all drugs, including the ADC treatment sacituzumab-govitecan (SG), with a higher effect on carboplatin and paclitaxel. In parallel, we established a dedicated pipeline in which freshly obtained biopsies were provided directly by radiologists (sample #6) during the procedure and immediately processed for integration into our biopsy-chips. Notably, the first fresh biopsy obtained yielded a high cellularity of approximately 10^6^ tumor cells (**Fig. S8B**), enabling robust on-chip drug testing. After exposure to standard-of-care agents, including three chemotherapies at 50 µM (5-FU, capecitabine, and paclitaxel) and two ADCs (SG and T-DXd), no significant reduction in cell viability at day 4 was observed (**Fig. 5G**). This highly chemoresistant profile mirrored the clinical course of the patient, who initially presented with a localized TNBC and demonstrated no response to neoadjuvant chemoimmunotherapy (anthracyclines, cyclophosphamide, carboplatin, paclitaxel, and pembrolizumab) with further disease progression under adjuvant capecitabine–pembrolizumab (59).

Altogether, these findings support the use of this *ex vivo* platform for rapid assessment of individualized drug responses under physiologically relevant conditions. Its successful application to primary tumor samples and biopsies underscores its translational relevance for predictive applications in clinics.

## Discussion

In this study, we report the use of a tumor-on-chip platform for rapid *ex vivo* drug testing in patient-derived breast cancer cells. Drug sensitivity profiles identified using this system closely mirrored the responses observed in matched PDX models *in vivo*, even when initiated from minimal biopsy-derived cell input. Therefore, we applied the ToC assay to surgical specimens and patient biopsies, where the observed sensitivity profiles reflected patients’ recent clinical responses, suggesting its potential as a practical and scalable tool for personalized treatment guidance.

We first demonstrated that the ToC platform is a reliable model, achieving an establishment rate – *i*.*e*., the number of samples that present more than 50% of cell viability onchip at day 0 – of 99% (n=86 ToC models established out of 88 PDX tumors, 5 out of 5 primary tumors, and 1 out of 1 patient biopsy). On average, the PDX-tissue from one mouse provides approximately 10^6^ cancer cells, sufficient to produce up to 25 chips containing 4×10^4^ cells each. This allows the evaluation of up to five drugs with three technical replicates per condition, whereas conventional *in vivo* testing requires one mouse per condition. Technical replicates exhibited low inter-assay variability and consistent drug response patterns across independent experiments, underscoring the robustness and high reproducibility of the ToC system. With this technique, we were able to discriminate sensitivity and resistance profiles in ToC. We showed concordance with matched *in vivo* responses in PDXs across six models of different molecular subtypes. Therefore, by drastically increasing the number of assays while using fewer cancer cells and reducing the turnaround time from weeks for PDOs and months for PDXs (60) to just 4 days, ToC offers a rapid, actionable, and reproducible platform that can be easily integrated into clinical workflows.

Although PDX tissue provides abundant cellular material, clinical samples typically rely on biopsies with limited cell numbers. In commercially available microfluidic chips, more than one-third of the input cells are lost within the dead volume of the device – a constraint incompatible with clinical applications. To address this limitation and maximize the use of scarce patient-derived material, we engineered a new microfluidic chip specifically designed for low-input samples. This custom-chip features an experimental working volume of 100% and a circular architecture designed to support further automation and high-throughput processing. Using biopsy material, we successfully established functional ToC models derived from both PDX-derived biopsies and primary human breast cancer samples, including one biopsy, thereby providing a proof-of-concept for the platform’s applicability to clinically relevant tumor specimens, thus supporting rapid personalized therapeutic decision-making. To overcome the intratumoral heterogeneity, such platforms can be multiplexed to several biopsies of the same patient, offering an internal and accurate comparison for each patient.

Beyond mirroring PDX profiles, ToC assays, from biopsy or resection samples, importantly reflect the histories and clinical outcomes of the original patients. The HBCx-14 model, derived from a chemo-naïve localized TNBC, showed broad chemosensitivity in ToC, whereas HBCx-239, originating from a heavily pre-treated metastatic ER+ BC, exhibited high chemoresistance on-chip. The latter patient did not respond to neoadjuvant taxane-based chemotherapy or adjuvant capecitabine. Consistently, neither the corresponding PDX model nor the matched ToC platform showed sensitivity to paclitaxel or 5FU, further supporting the concordance between clinical outcomes, *in vivo* PDX responses, and *ex vivo* ToC sensitivity. However, both ToC and PDX display a few discrepancies with patient tumor progression: HBCx-73 showed sensitivity to 5FU in both PDX and ToC, whereas the disease progressed in patients; conversely, HBCx-178 displayed stable disease in patients but was classified as resistant in both models. To discriminate drug sensitivity in ToC, we have considered a 33% threshold based on the corresponding PDX tumor response to drug, but it may thus not fully capture the spectrum of drug responses *in vivo*, particularly subtle cytostatic effects. Since patient survival remains the most critical and relevant readout for these functional assays, this 33% threshold must be, in the future, carefully compared with survival data, and finer efficacy categories of drug efficacy in ToC may be defined through clinical assays in large patient cohorts.

Another major challenge for ToC assays is ensuring that drug concentrations applied *ex vivo* accurately reflect pharmacologically relevant exposures achieved in patients *in vivo*. In this study, drug concentrations were selected based on pharmacokinetic data from prior clinical studies (54) and applied uniformly across all PDX models. While this strategy ensured experimental comparability, it may have limited translational validity in tumor models. The only false-negative ToC result involved 5FU at 50 µM (HBCx-178), and increasing 5FU concentration up to 400 µM (54) restored sensitivity. Conversely, the only false-positive result related to HBCx-157, known to be resistant to paclitaxel *in vivo* and classified as sensitive in the ToC assay. This misclassification occurred at drug concentrations exceeding tenfold the maximum plasma levels reported in patients (54), and the ToC assay correctly identified this model as resistant for physiologically relevant concentrations (between 1 and 10 µM), while consistently identifying HBCx-14 and HBCx-164 models as sensitive and HBCx-178, HBCx-239, and HBCx-73 models as resistant in these concentrations. Collectively, these findings highlight the assay’s high specificity in ToC and, importantly, support using pharmacologically achievable plasma levels for comparison with patient outcomes, ensuring predictive accuracy while avoiding clinically irrelevant supra- or infra-physiological exposures.

Finally, to mirror more recent clinical practice, the antibody–drug conjugates (ADC), which combine targeted therapy and chemotherapy, currently represent a turnaround in anti-cancer treatment and have been tested in ToC. T-DXd, composed of a humanized anti-HER2 IgG1 monoclonal antibody linked to a potent topoisomerase I inhibitor (DXd) via a cleavable linker (61), was evaluated on-chip in two HER2-positive models: HBCx-239 (HER2 1+) and HBCx-73 (HER2 3+); and one HER2-negative model (HER2 0), the HBCx-264. The HER2 models showed high resistance to T-DXd, while in both HER2-positive tumors, increasing concentrations led to enhanced cytotoxicity. However, the addition of aphidicolin, an inhibitor of DNA polymerase, decreases such cytotoxicity, revealing the need for active DNA replication to maximize payload-induced cytotoxicity, a key determinant of T-DXd efficacy. Other mechanisms of T-DXd, including HER2 binding and Fc γ RIII-driven immune engagement, could also be dissected with ToC platforms. For instance, cathepsin, recently identified as a protease mediating linker cleavage and payload release in HER2-low tumors (62), can be selectively incorporated on-chip to probe its functional role. Moreover, in our current PDX-derived models, the absence of immune effector cells may have led to an underestimation of T-DXd activity, emphasizing the need to integrate immune components in future assays. This is particularly relevant for T-DXd, where Fc-mediated mechanisms such as ADCC contribute to therapeutic efficacy (63). Therefore, the growing ability of ToC systems to support autologous stromal and immune co-cultures provides the opportunity to reconstruct tumor-specific immune contexts and capture the full spectrum of ADC and immunotherapy activity (38, 39), especially in the context of personalized therapeutic decisions, which require performing experiments in the most relevant and physiological cellular microenvironment.

To conclude, we show that ToC models offer a powerful and complementary alternative to patient-derived avatars such as PDXs and PDOs, enabling rapid and individualized preclinical testing with potential for large-scale integration into breast cancer care. Looking forward, collective efforts will be critical to validate the TOC platform in a larger cohort of tumors and treatments, and to establish their clinical validity by correlating on-chip drug responses from predefined conditions with patient outcomes. Thereby, this will position ToC platforms as next-generation functional avatars for precision oncology.

## Acknowledgments

This project has been supported by ITMO Cancer (23CP060-00) of Aviesan within the framework of the 2021-2030 Cancer Control Strategy (Inserm), the Agence Nationale de Recherche (ANR-24-EXME-0005 - PEPR MEDOOC - TME-on-Chip), the Institute of Women’s Cancer IHU (ANR23-IAHU-0006), Fondation de France (00149014/WB-2023-50799), the Horizon Europe programme (101136464), and Viasanté Mutuelle. This work also benefited from the technical contribution of the joint service unit IPGG Technological Platform CNRS UAR 3750. We would like to thank the engineers of this unit, in particular Lars Kool and Elian Martin, for their advice during the development of the experiments.

## Author contributions

CH, JJ, NS, and AV performed the ToC experiments. MN performed the live experiments in ToC. EM, LS, HD, AB and EM performed the PDX experiments. LP analysed the data. MS helped design the analyses. CH and LP created the figures and wrote the manuscript. GZ, EM, LC, MCP, and SD designed the study and revised the manuscript. AG, AVS, TR and LC set the clinical pipeline and supplied human samples. MCP and SD secured funding.

## Materials and Methods

### PDX establishment and *in vivo* efficacies

PDX models of BC were generated at Institut Curie (LIP, Paris, France) using specimens from standard-of-care surgical resections or biopsies of metastatic sites from patients with BC as previously detailed (64, 65). Non-necrotic tumor fragments of 30-60 mm^3^ were heterotopically implanted into the interscapular fat pad of female Swiss nude immunocompromised mice (Charles River Laboratories). The experiments were conducted following institutional guidelines and the rules of the French Ethics Committee: CEEA-IC (Comité d’Ethique en matière d’expérimentation animale de l’Institut Curie, National registration number: #118); project authorization no.: APAFIS#25870-2020060410487032. Once established, PDXs were histologically characterized to confirm similarity to the corresponding patient tumor. Several molecular subtypes of BC were engrafted to capture the disease’s heterogeneity: four triple-negative breast cancers (TNBC, HBCx-14, HBCx-157, HBCx-164, and HBCx-178), one HER2 3+ (HBCx-73), and one ER+ (HBCx-239) (65–67). For each BC subtype, PDX were then divided in two groups: one group not treated (control) and one group in which mice were treated with at least 3 clinically relevant drugs: carboplatin (alkylating agent, 90mg/kg, one intraperitoneal injection every 3 weeks) (65), capecitabine (anti-metabolite, oral prodrug of 5FU, 540 mg/kg, 5 times a week per os) and paclitaxel (anti-microtubule agent, 25 mg/kg, one intraperitoneal injection once a week). Cisplatin (alkylating agent, 6mg/kg, one intraperitoneal injection every 3 weeks) was also tested on the HBCx-14 model (64). Additionally, trastuzumab-deruxtecan (an antibody-drug conjugate targeting HER2, 10 mg/kg or 4 mg/kg one intraperitoneal injection at D1) was tested on HBCx-73 (HER2 3+) and HBCx-239 (ER+ HER2 1+), respectively. Tumor growth was evaluated twice weekly by measurement of two perpendicular diameters of tumors using calipers over 40 days or until ethical size was reached (2000 mm^3^), whichever occurred first. Tumor volume was approximated as

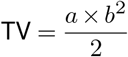

where *a* and *b* represent the longest and shortest diameters, respectively. For each tumor, volumes were reported to the initial volume as relative tumor volume (RTV). Means (and SE) of RTV in the same treatment group were calculated, and growth curves were established as a function of time. Tumor growth inhibition (TGI) was assessed by comparing mean tumor volumes in the treated and control groups at the same time point, using the formula:

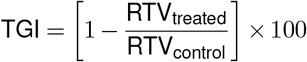

and changes in tumor volume were calculated in the treated groups as follows:

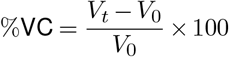

where *V*_0_ is the tumor volume at baseline and *V*_*t*_ is the tumor volume at the indicated time point (23, 52). PDX models were classified as either sensitive or resistant to treatment *in vivo* based on the TGI and %VC. Tumors were considered sensitive when the TGI was above 90% and resistant below 60%. For intermediate cases (60% *<* TGI *<* 90%), classification was refined using the %VC in the treated group and tumors were categorized as resistant if they displayed progressive disease (%VC *>* +35%), and as sensitive if they exhibited either stable disease (− 50% ≤ %VC ≤ +35%) or tumor response (%VC ≤ −50%) (52).

### Tumor dissociation of breast cancer PDX

Tumors from the control groups of PDX models were resected and sent immediately to MMBM laboratory at room temperature in a 50 mL Falcon filled with DMEM. The PDX tumors measured approximately 20 × 15 mm (*±* 2 mm) upon collection, corresponding to the final tumor volume observed in the *in vivo* experiments. Tumor tissues were dissociated into single-cell suspensions by combining mechanical dissociation with enzymatic degradation of the extracellular matrix to maintain the structural integrity of tissues. The protocol used was compatible for both primary human tumor tissues and xenografts, and optimized for a wide range of tumor types, including BC. Directly upon arrival, tumors were manually minced into small fragments (1-2 mm^3^) with a scalpel and enzymatically digested with a tumor dissociation kit (Miltenyi Biotec, #130-095-929), according to the manufacturer’s instructions, and placed in a thermomixer (37°C, 500 RPM) for 45 minutes. After dissociation, samples were applied to a filter (70 µm strainer) to remove any remaining larger particles from cell suspensions and placed in medium A: RPMI-1640 (GE healthcare) supplemented with 10% human AB serum (Institut Jacques Boy, Reims, France), 1% of Sodium Pyruvate, and 0.1% Penicillin/Streptomycin before proceeding to tumoral cells isolation.

The magnetic cell sorting (MACS) (Miltenyi, 130-091-051) was used along with the requisite kits in order to isolate the tumor cells from the different TME components. First, red blood cells were lysed using the Red Blood Cell Lysis Solution (Miltenyi Biotec, #130-094-183), which is suitable for single-cell suspensions of both human and mouse origin. Then, cell debris which often occurs after tissue dissociation and impairs downstream applications were removed using a density gradient and the Debris Removal Solution (Miltenyi Biotec, #130-109-398). Third, dead cells were magnetically labelled using the Dead Cell Removal Microbeads (Miltenyi Biotec, #130-090-101) and passed through a magnetic separation column according to the supplier’s protocol. Since microbeads target phosphatidylserine, early apoptotic cells with an intact cellular membrane were also retained in the column and therefore eliminated from the samples. Finally, mouse cells were magnetically labelled with a cocktail of monoclonal antibodies conjugated with MACS Microbeads (Miltenyi Biotec, #130-104-694) and the pellet was placed in the magnetic field of the separation column. Mouse cells were retained within the column, whereas the unlabelled viable human tumoral cells that ran through the column were collected, counted, and processed for the tumor-on-chip preparation (see below).

### Breast-cancer patient samples

Fresh human tumor samples were provided by the Pathology Department of Institut Curie (Paris, France) from patients who received a standard-of-care surgical resection of breast cancer (patients #1, #2, #4). Tissue specimens corresponded to surgical residues available after histopathological examination and deemed unnecessary for diagnostic purposes. Tumor samples were dissociated using an adapted and simplified version of the protocol previously described in the BC PDX tumor dissociation section. For the preparation of human-derived ToC, only red blood cells and debris were removed using the same techniques and reagents as described above. We subsequently implemented a dedicated pipeline to allow direct processing of freshly obtained biopsies, which were directly transferred to our laboratory by the radiology department, for integration into the ToC platform (patient #3). Patients were informed of the potential use of their samples and related data for research, and they provided a written non-opposition form according to the French rules on non-interventional studies using leftover biological specimens for research. The human experimental procedures follow the Declaration of Helsinki guidelines.

### Tumor-on-Chip preparation

Microfluidic devices (#DAX-1) were purchased from AIM-Biotech. A matrix composed of type I rat tail collagen (Thermofisher, #A1048301) was prepared at the final concentration of 2.5 mg/mL according to manufacturer protocol with 10X PBS (Gibco, #12579099), distilled H_2_O (Gibco, #15230162) and pH neutralisation of the resulting collagen mixture to 7.0 using 1N NaOH solution (Sigma-Aldrich, #S2770). Following tumor dissociation, tumor cells were counted, suspended in the freshly prepared collagen matrix, and manually injected into the central chamber of the DAX-1 chips at a final density of 4000 tumor cells/µL. All pipettes, tips, chips, and reagents were kept chilled to pre-empt premature polymerization of the gel. Each chip was loaded with 10 µL of gel equivalent to 4×10^4^ tumor cells/chip. After the addition of the gel in the microfluidic device, the chip was incubated in a humidified chamber for 50 min in the incubator (37 °C, 5% CO_2_). After gel polymerization, 200 µL of either medium A alone (control) or containing increasing concentrations of drugs was added to the lateral chambers of the chip: carboplatin, cisplatin, 5FU, paclitaxel, trastuzumab-deruxtecan. Drugs were given by the Institut Curie pharmacy. For each BC-PDX model, at least three biological replicates were performed (*i*.*e*., the same breast cancer model was engrafted in three mice or more); and for each biological replicate, at least two technical replicates (*i*.*e*., two collagen gels containing tumor cells isolated from the same PDX tumor) were tested and analyzed for each drug concentration.

### Drug sensitivity assay

Tumor cell viability was quantified after 96h of drug exposure on the ToC platform using the LIVE/DEAD™ Viability/Cytotoxicity Kit (ThermoFisher, #R37601). Live cells were stained with calcein-AM (GFP), whereas dead cells were labelled with ethidium homodimer-1 (Cy5). Images were acquired with a spinning disk Leica, Objective 10X, NA 0.3.

The LIVE/DEAD™ assay was performed every 24h from D0 to D4, to both determine the optimal time point for drug sensitivity assessment and to characterize response kinetics. For each drug, cell viability was assessed following exposure to a harmonized panel of four concentrations (1, 10, 20, and 50 µM), except for trastuzumab-deruxtecan, which was tested at 0.01, 0.1, 0.5, 1, 10, 100, and 1000 µg/mL. Viability values were normalized to both the untreated control and the baseline measurement at D0. For aphidicolin experiments, following gel polymerization, cells were pre-incubated for 5 minutes at 37°C with medium A supplemented with aphidicolin (2.95 µM), prior to the addition of either medium A alone or T-DXd into the lateral chambers of the chips. For patient samples, a fixable Image-iT DEAD Green Viability Stain kit (Ther-moFisher, #I10291) was used to selectively label dead cells with a live-cell impermeant DNA-binding dye (GFP). Chips were fixed with 4% PFA for 20 minutes, permeabilized with PBS containing 0.1% Triton for 2 hours, and stained overnight with anti-pancytokeratin antibody (Cy3, ThermoFisher, #42-9003-82). Nuclei were counterstained with Hoechst to enable total cell counting. Human cancer cells were identified as pancytokeratin-positive, and viability was quantified based on the DEAD Green signal.

### Image analysis

For each chip, nine fields of view were acquired using a 10X objective. For PDX-derived samples, live and dead cell counts were obtained using Meta-Morph® and ImageJ. Viability was calculated for each field as the percentage of live cells, then averaged across the nine images per chip. The resulting mean viability was normalized to the D0 control (measured immediately after seeding in collagen). For human tumor samples, dead tumor cells — identified as pancytokeratin-positive and stained green — were quantified using ImageJ (Version 2.16.0/1.54p). Tumor cell death was expressed as the percentage of dead tumor cells, averaged across the nine fields per chip, and normalized to the D0 control. These values were then compared to the corresponding untreated control condition.

### Microfluidic setup for live cell imaging

To validate our endpoint measurement method, we compared it with the cell-death dynamics observed by live-cell imaging the chips. An adapted version of our previously described microfluidic setup (39) was used to continuously image cells within the microfluidic device. Briefly, the tumor-on-chips were mounted on the motorized stage of an inverted wide-field fluorescence video-microscope (DMi8, Leica and TI3, Nikon), enclosed in an incubation chamber to provide a humid atmosphere with 5% CO_2_ at 37°C. Images were taken by using either a Photometrics Prime BSI Express camera and a CoolLed PE4000 light source (Nikon TI3) or a Retiga R6 camera and Lumencor SOLA SE 365 light engine (Leica DMi8). All acquisitions were carried out using the Meta-Morph software. A 5X objective was used. For each gel, 2-4 positions were acquired. For cancer death analysis, 6 mM of CellEvent Caspase-3/7 Green Detection Reagent (Thermofisher, #C10423, green fluorescence) was added to the culture medium, and image acquisition was carried out for each position every 30–60 min for 48-96h. Resulting analysis of apoptotic signaling on acquired video was carried out using ImageJ and TM-STAMP software as described in our prior publications (39).

### Establishment of Biopsy-on-Chip

To simulate a clinical biopsy, PDX and patient tumor samples were biopsied *ex vivo* immediately upon arrival in the laboratory. An 18-gauge core needle, corresponding to a diameter of ≥ 1.02 mm, was passed three or four times through each tissue specimen, depending on the maximal length of the sample. Specimens were biopsied three times when the maximum length was 20 mm and four times when the maximum length was < 20 mm. This sampling strategy was established in close collaboration with radiologists, aiming to balance the need for sufficient cellular input for down-stream assays with the practical constraints of routine clinical procedures. For each tumor, the resulting biopsy cores were collected in a separate Eppendorf tube, while the remaining tumor tissue was minced into fragments (1-2 mm^3^) using a scalpel. Both sample types, biopsy cores and whole-tumor fragments, were then dissociated in par-allel with the same dissociation kits and protocols described above (Miltenyi Biotec, #130-091-051, #130-094-183, #130-109-398) to isolate tumoral cells and proceed to the biopsy-on-chip preparation (see below).

### Biopsy-on-Chip preparation: Microfluidic Device Design and Fabrication

We engineered a custom microfluidic device with no dead volume and a minimized volume of interest, specifically optimized for high-resolution imaging. In contrast, commercially available platforms such as the AIM Biotech (#DAX-1) chip allow imaging of less than one-third of the device’s total area. Indeed, a single 10x objective field covers only 0.479 mm^2^, approximately 1/28th of the chip’s available imaging surface (excluding depth), leaving a substantial fraction of the embedded cells unanalyzed. This inefficiency becomes particularly critical when working with the limited number of tumor cells obtained from small tissue samples such as core needle biopsies. To overcome this constraint, our device was specifically designed to maximize the proportion of analyzable cells, thereby enhancing both experimental throughput and imaging fidelity. For rapid prototyping and precise customization of channel geometry, molds were made from stiff plastic using a Mini-Mill/GX CNC micromilling machine (Minitech Machinery Corporation) from the IPGG UAR 3750 technology platform. The mold was then sent to our industrial partner for assembly into a steel mold frame. The molded chips were then returned to IPGG for drilling and were assembled with a 60 µm-thick COC-6013 film using laser welding (Turn-Key S system, Probylas). The 3D-printed molds were subsequently exposed to vapor phase silanization using trichloro(1H,1H,2H,2H-perfluorooctyl)silane (Sigma-Aldrich, #448931) to prevent liquid leakage and improve PDMS demolding quality. To eliminate potential cytotoxic effects from residual silane, the molds were thoroughly rinsed with isopropanol before use. Chips were fabricated by casting PDMS (Sylgard 184, Farnell, #101697) at a 10:1 w/w ratio of elastomer to curing agent. The mixture was degassed in a desiccator to remove air bubbles, then cured at 75°C overnight. Once cured, the PDMS was peeled from the mold, and inlets of 1 mm diameter were created with a 1 mm puncher (World Precision Instrument, #15453532). Devices were cleaned with isopropanol, dried with compressed air, and residual particles were removed from the PDMS surface using transparent adhesive tape (Scotch Tape, 3M). The chips were then irreversibly bonded to 1.7 mm–thick glass coverslips via oxygen plasma treatment.

### Statistical analysis

Wilcoxon’s tests in R software (RStudio Version 2024.12.1+563 and R Version 4.4.2) were used for statistical analysis. A minimum of two biological replicates were used for each breast cancer model, with at least two technical replicates for each condition, and the standard deviation is presented as error bars.

## References

1. Freddie Bray, Mathieu Laversanne, Hyuna Sung, Jacques Ferlay, Rebecca L. Siegel, Isabelle Soerjomataram, and Ahmedin Jemal. Global cancer statistics 2022: Globocan estimates of incidence and mortality worldwide for 36 cancers in 185 countries. CA: A Cancer Journal for Clinicians, 74(3): 229–263, 2024. doi: 10.3322/caac.21834.

2. Joanne Kim, Andrew Harper, Valerie McCormack, Hyuna Sung, Nehmat Houssami, Eileen Morgan, Miriam Mutebi, Gail Garvey, Isabelle Soerjomataram, and Miranda M. Fidler-Benaoudia. Global patterns and trends in breast cancer incidence and mortality across 185 countries. Nature Medicine, 31(4):1154–1162, Apr 2025. ISSN 1546-170X. doi: 10.1038/s41591-025-03502-3.

3. Kimberly H. Allison, M. Elizabeth H. Hammond, Mitchell Dowsett, Shannon E. McKernin, Lisa A. Carey, Patrick L. Fitzgibbons, Daniel F. Hayes, Sunil R. Lakhani, Mariana Chavez-MacGregor, Jane Perlmutter, Charles M. Perou, Meredith M. Regan, David L. Rimm, W. Fraser Symmans, Emina E. Torlakovic, Leticia Varella, Giuseppe Viale, Tracey F. Weisberg, Lisa M. McShane, and Antonio C. Wolff. Estrogen and progesterone receptor testing in breast cancer: Asco/cap guideline update. Journal of Clinical Oncology, 38(12):1346–1366, 2020. doi: 10.1200/JCO.19.02309. PMID: 31928404.

4. Harold J. Burstein, Angela DeMichele, Lesley Fallowfield, Mark R. Somerfield, N. Lynn Henry, null null, N. Lynn Henry, Zoneddy Dayao, Anthony Elias, Kevin Kalinsky, Lisa M. McShane, Beverly Moy, Ben Ho Park, Kelly M. Shanahan, Priyanka Sharma, Rebecca Shatsky, Erica Stringer-Reasor, Melinda Telli, Nicholas C. Turner, Angela DeMichele, Harold J. Burstein, Debra L. Barton, Ali Dorris, Lesley J. Fallowfield, Dharamvir Jain, Stephen R.D. Johnston, Larissa A. Korde, Jennifer K. Litton, Erin R. Macrae, Lindsay L. Peterson, Praveen Vikas, Rachel L. Yung, and Hope S. Rugo. Endocrine and targeted therapy for hormone receptor–positive, human epidermal growth factor receptor 2–negative metastatic breast cancer—capivasertib-fulvestrant: Asco rapid recommendation update. Journal of Clinical Oncology, 42(12): 1450–1453, 2024. doi: 10.1200/JCO.24.00248. PMID: 38478799.

5. Elisa Agostinetto, Giuseppe Curigliano, and Martine Piccart. Emerging treatments in her2-positive advanced breast cancer: Keep raising the bar. Cell Reports Medicine, 5(6): 101575, 2024. ISSN 2666-3791. doi: 10.1016/j.xcrm.2024.101575.

6. Carsten Denkert, Cornelia Liedtke, Andrew Tutt, and Gunter von Minckwitz. Molecular alterations in triple-negative breast cancer—the road to new treatment strategies. The Lancet, 389(10087): 2430–2442, 2017. ISSN 0140-6736. doi: 10.1016/S0140-6736(16)32454-0.

7. Eduarda Carvalho, Sule Canberk, Fernando Schmitt, and Nuno Vale. Molecular subtypes and mechanisms of breast cancer: Precision medicine approaches for targeted therapies. Cancers, 17(7), 2025. ISSN 2072-6694. doi: 10.3390/cancers17071102.

8. K. Hanna and K. Mayden. Chemotherapy treatment considerations in metastatic breast cancer. JADPRO, 12(Suppl 2):6–12, 2021. doi: 10.6004/jadpro.2021.12.2.11. Epub 2021 Mar 1.

9. Christophe Le Tourneau, Jean-Pierre Delord, Anthony Gonçalves, Céline Gavoille, Coraline Dubot, Nicolas Isambert, Mario Campone, Olivier Trédan, Marie-Ange Massiani, Cécile Mauborgne, Sebastien Armanet, Nicolas Servant, Ivan Bièche, Virginie Bernard, David Gentien, Pascal Jezequel, Valéry Attignon, Sandrine Boyault, Anne Vincent-Salomon, Vincent Servois, Marie-Paule Sablin, Maud Kamal, and Xavier Paoletti. Molecularly targeted therapy based on tumour molecular profiling versus conventional therapy for advanced cancer (shiva): a multicentre, open-label, proof-of-concept, randomised, controlled phase 2 trial. The Lancet Oncology, 16(13): 1324–1334, 2015. ISSN 1470-2045. doi: 10.1016/S1470-2045(15)00188-6.

10. Christophe Massard, Stefan Michiels, Charles Ferté, Marie-Cécile Le Deley, Ludovic Lacroix, Antoine Hollebecque, Loic Verlingue, Ecaterina Ileana, Silvia Rosellini, Samy Ammari, Maud Ngo-Camus, Rastislav Bahleda, Anas Gazzah, Andrea Varga, Sophie Postel-Vinay, Yohann Loriot, Caroline Even, Ingrid Breuskin, Nathalie Auger, Bastien Job, Thierry De Baere, Frederic Deschamps, Philippe Vielh, Jean-Yves Scoazec, Vladimir Lazar, Catherine Richon, Vincent Ribrag, Eric Deutsch, Eric Angevin, Gilles Vassal, Alexander Eggermont, Fabrice André, and Jean-Charles Soria. High-throughput genomics and clinical outcome in hard-to-treat advanced cancers: Results of the moscato 01 trial. Cancer Discovery, 7(6):586–595, Jun 2017. ISSN 2159-8274. doi: 10.1158/2159-8290.CD-16-1396.

11. Jason K. Sicklick, Shumei Kato, Ryosuke Okamura, Maria Schwaederle, Michael E. Hahn, Casey B. Williams, Pradip De, Amy Krie, David E. Piccioni, Vincent A. Miller, Jeffrey S. Ross, Adam Benson, Jennifer Webster, Philip J. Stephens, J. Jack Lee, Paul T. Fanta, Scott M. Lippman, Brian Leyland-Jones, and Razelle Kurzrock. Molecular profiling of cancer patients enables personalized combination therapy: the i-predict study. Nature Medicine, 25(5):744– 750, May 2019. ISSN 1546-170X. doi: 10.1038/s41591-019-0407-5.

12. Alice P. Chen, Shivaani Kummar, Nancy Moore, Lawrence V. Rubinstein, Yingdong Zhao, P. Mickey Williams, Alida Palmisano, David Sims, Geraldine O’Sullivan Coyne, Christina L. Rosenberger, Mel Simpson, Kanwal P. S. Raghav, Funda Meric-Bernstam, Stephen Leong, Saiama Waqar, Jared C. Foster, Mariam M. Konaté, Biswajit Das, Chris Karlovich, Chih-Jian Lih, Eric Polley, Richard Simon, Ming-Chung Li, Richard Piekarz, and James H. Doroshow. Molecular profiling-based assignment of cancer therapy (nci-mpact): A randomized multicenter phase ii trial. JCO Precision Oncology, (5):133–144, 2021. doi: 10.1200/PO.20.00372. PMID:.

13. Funda Meric-Bernstam, James M. Ford, Peter J. O’Dwyer, Geoffrey I. Shapiro, Lisa M. McShane, Boris Freidlin, Roisin E. O’Cearbhaill, Suzanne George, Julia Glade-Bender, Gary H. Lyman, James V. Tricoli, David Patton, Stanley R. Hamilton, Robert J. Gray, Douglas S. Hawkins, Bhanumati Ramineni, Keith T. Flaherty, Petros Grivas, Timothy A. Yap, Jordan Berlin, James H. Doroshow, Lyndsay N. Harris, Jeffrey A. Moscow, and on behalf of ComboMATCH study team. National cancer institute combination therapy platform trial with molecular analysis for therapy choice (combomatch). Clinical Cancer Research, 29(8): 1412–1422, Apr 2023. ISSN 1078-0432. doi: 10.1158/1078-0432.CCR-22-3334.

14. Ben Kinnersley, Amit Sud, Andrew Everall, Alex J. Cornish, Daniel Chubb, Richard Culliford, Andreas J. Gruber, Adrian Lärkeryd, Costas Mitsopoulos, David Wedge, and Richard Houlston. Analysis of 10,478 cancer genomes identifies candidate driver genes and opportunities for precision oncology. Nature Genetics, 56(9):1868–1877, Sep 2024. ISSN 1546-1718. doi: 10.1038/s41588-024-01785-9.

15. Jingwen Bai, Yiyang Gao, and Guojun Zhang. The treatment of breast cancer in the era of precision medicine. Cancer Biology & Medicine, 2025. ISSN 2095-3941. doi: 10.20892/j.issn.2095-3941.2024.0510.

16. Michele Masucci, Claes Karlsson, Lennart Blomqvist, and Ingemar Ernberg. Bridging the divide: A review on the implementation of personalized cancer medicine. Journal of Personalized Medicine, 14(6), 2024. ISSN 2075-4426. doi: 10.3390/jpm14060561.

17. Shahrokh Abdolahi, Zeinab Ghazvinian, Samad Muhammadnejad, Mahshid Saleh, Hamid Asadzadeh Aghdaei, and Kaveh Baghaei. Patient-derived xenograft (pdx) models, applications and challenges in cancer research. Journal of Translational Medicine, 20(1):206, May 2022. ISSN 1479-5876. doi: 10.1186/s12967-022-03405-8.

18. Yixin Shi, Zhanwen Guan, Gengxi Cai, Yichu Nie, Chuling Zhang, Wei Luo, and Jia Liu. Patient-derived organoids: a promising tool for breast cancer research. Frontiers in Oncology, Volume 14 - 2024, 2024. ISSN 2234-943X. doi: 10.3389/fonc.2024.1350935.

19. Anthony Letai, Patrick Bhola, and Alana L. Welm. Functional precision oncology: Testing tumors with drugs to identify vulnerabilities and novel combinations. Cancer Cell, 40(1): 26–35, 2022. ISSN 1535-6108. doi: 10.1016/j.ccell.2021.12.004.

20. Yoko S. DeRose, Guoying Wang, Yi-Chun Lin, Philip S. Bernard, Saundra S. Buys, Mark T. W. Ebbert, Rachel Factor, Cindy Matsen, Brett A. Milash, Edward Nelson, Leigh Neumayer, R. Lor Randall, Inge J. Stijleman, Bryan E. Welm, and Alana L. Welm. Tumor grafts derived from women with breast cancer authentically reflect tumor pathology, growth, metastasis and disease outcomes. Nature Medicine, 17(11):1514–1520, Nov 2011. ISSN 1546-170X. doi: 10.1038/nm.2454.

21. Yihan Liu, Wantao Wu, Changjing Cai, Hao Zhang, Hong Shen, and Ying Han. Patientderived xenograft models in cancer therapy: technologies and applications. Signal Transduction and Targeted Therapy, 8(1):160, Apr 2023. ISSN 2059-3635. doi: 10.1038/s41392-023-01419-2.

22. P. Cottu, E. Marangoni, F. Assayag, P. de Cremoux, A. Vincent-Salomon, Ch. Guyader, L. de Plater, C. Elbaz, N. Karboul, J. J. Fontaine, S. Chateau-Joubert, P. Boudou-Rouquette, S. Alran, V. Dangles-Marie, D. Gentien, M.-F. Poupon, and D. Decaudin. Modeling of response to endocrine therapy in a panel of human luminal breast cancer xenografts. Breast Cancer Research and Treatment, 133(2):595–606, Jun 2012. ISSN 1573-7217. doi: 10.1007/s10549-011-1815-5.

23. Elisabetta Marangoni, Anne Vincent-Salomon, Nathalie Auger, Armelle Degeorges, Franck Assayag, Patricia de Cremoux, Ludmilla de Plater, Charlotte Guyader, Gonzague De Pinieux, Jean-Gabriel Judde, Magali Rebucci, Carine Tran-Perennou, Xavier Sastre-Garau, Brigitte Sigal-Zafrani, Olivier Delattre, Veronique Dieras, and Marie-France Poupon. A new model of patient tumor-derived breast cancer xenografts for preclinical assays. Clinical Cancer Research, 13(13):3989–3998, Jul 2007. ISSN 1078-0432. doi: 10.1158/1078-0432.CCR-07-0078.

24. Xiaomei Zhang, Sofie Claerhout, Aleix Prat, Lacey E. Dobrolecki, Ivana Petrovic, Qing Lai, Melissa D. Landis, Lisa Wiechmann, Rachel Schiff, Mario Giuliano, Helen Wong, Suzanne W. Fuqua, Alejandro Contreras, Carolina Gutierrez, Jian Huang, Sufeng Mao, Anne C. Pavlick, Amber M. Froehlich, Meng-Fen Wu, Anna Tsimelzon, Susan G. Hilsenbeck, Edward S. Chen, Pavel Zuloaga, Chad A. Shaw, Mothaffar F. Rimawi, Charles M. Perou, Gordon B. Mills, Jenny C. Chang, and Michael T. Lewis. A renewable tissue resource of phenotypically stable, biologically and ethnically diverse, patient-derived human breast cancer xenograft models. Cancer Research, 73(15):4885–4897, Jul 2013. ISSN 0008-5472. doi: 10.1158/0008-5472.CAN-12-4081.

25. Peter Kabos, Jessica Finlay-Schultz, Chunling Li, Enos Kline, Christina Finlayson, Joshua Wisell, Christopher A. Manuel, Susan M. Edgerton, J. Chuck Harrell, Anthony Elias, and Carol A. Sartorius. Patient-derived luminal breast cancer xenografts retain hormone receptor heterogeneity and help define unique estrogen-dependent gene signatures. Breast Cancer Research and Treatment, 135(2):415–432, Sep 2012. ISSN 1573-7217. doi: 10.1007/s10549-012-2164-8.

26. Le Tong, Weiyingqi Cui, Boya Zhang, Pedro Fonseca, Qian Zhao, Ping Zhang, Beibei Xu, Qisi Zhang, Zhen Li, Brinton Seashore-Ludlow, Ying Yang, Longlong Si, and Andreas Lundqvist. Patient-derived organoids in precision cancer medicine. Med, 5(11): 1351–1377, 2024. ISSN 2666-6340. doi: 10.1016/j.medj.2024.08.010.

27. Soon-Chan Kim, Ji Won Park, Ha-Young Seo, Minjung Kim, Jae-Hyeon Park, Ga-Hye Kim, Ja Oh Lee, Young-Kyoung Shin, Jeong Mo Bae, Bon-Kyoung Koo, Seung-Yong Jeong, and Ja-Lok Ku. Multifocal organoid capturing of colon cancer reveals pervasive intratumoral heterogenous drug responses. Advanced Science, 9(5): 2103360, 2022. doi: 10.1002/advs.202103360.

28. Jérôme Cartry, Sabrina Bedja, Alice Boilève, Jacques R. R. Mathieu, Emilie Gontran, Maxime Annereau, Bastien Job, Ali Mouawia, Pierre Mathias, Thierry D. Baère, Antoine Italiano, Benjamin Besse, Isabelle Sourrouille, Maximiliano Gelli, Mohamed-Amine Bani, Peggy Dartigues, Antoine Hollebecque, Cristina Smolenschi, Michel Ducreux, David Malka, and Fanny Jaulin. Implementing patient derived organoids in functional precision medicine for patients with advanced colorectal cancer. Journal of Experimental & Clinical Cancer Research, 42(1):281, Oct 2023. ISSN 1756-9966. doi: 10.1186/s13046-023-02853-4.

29. G. Emerens Wensink, Sjoerd G. Elias, Jasper Mullenders, Miriam Koopman, Sylvia F. Boj, Onno W. Kranenburg, and Jeanine M. L. Roodhart. Patient-derived organoids as a predictive biomarker for treatment response in cancer patients. npj Precision Oncology, 5(1):30, Apr 2021. ISSN 2397-768X. doi: 10.1038/s41698-021-00168-1.

30. Alice Boilève, Jérôme Cartry, Negaar Goudarzi, Sabrina Bedja, Jacques R.R. Mathieu, Mohamed-Amine Bani, Rémy Nicolle, Ali Mouawia, Ryme Bouyakoub, Claudio Nicotra, Maud Ngo-Camus, Bastien Job, Karélia Lipson, Valérie Boige, Marine Valéry, Anthony Tarabay, Peggy Dartigues, Lambros Tselikas, Thierry de Baere, Antoine Italiano, Simona Cosconea, Maximiliano Gelli, Elena Fernandez-de Sevilla, Maxime Annereau, David Malka, Cristina Smolenschi, Michel Ducreux, Antoine Hollebecque, and Fanny Jaulin. Organoids for functional precision medicine in advanced pancreatic cancer. Gastroenterology, 167(5): 961–976.e13, Oct 2024. ISSN 0016-5085. doi: 10.1053/j.gastro.2024.05.032.

31. Raphaëlle Servant, Michele Garioni, Tatjana Vlajnic, Melanie Blind, Heike Pueschel, David C Müller, Tobias Zellweger, Arnoud J Templeton, Andrea Garofoli, Sina Maletti, Salvatore Piscuoglio, Mark A Rubin, Helge Seifert, Cyrill A Rentsch, Lukas Bubendorf, and Clémentine Le Magnen. Prostate cancer patient-derived organoids: detailed outcome from a prospective cohort of 81 clinical specimens. The Journal of Pathology, 254(5): 543–555, 2021. doi: 10.1002/path.5698.

32. Norman Sachs, Joep de Ligt, Oded Kopper, Ewa Gogola, Gergana Bounova, Fleur Weeber, Anjali Vanita Balgobind, Karin Wind, Ana Gracanin, Harry Begthel, Jeroen Korving, Ruben van Boxtel, Alexandra Alves Duarte, Daphne Lelieveld, Arne van Hoeck, Robert Frans Ernst, Francis Blokzijl, Isaac Johannes Nijman, Marlous Hoogstraat, Marieke van de Ven, David Anthony Egan, Vittoria Zinzalla, Jurgen Moll, Sylvia Fernandez Boj, Emile Eugene Voest, Lodewyk Wessels, Paul Joannes van Diest, Sven Rottenberg, Robert Gerhardus Jacob Vries, Edwin Cuppen, and Hans Clevers. A living biobank of breast cancer organoids captures disease heterogeneity. Cell, 172(1):373–386.e10, 2018. ISSN 0092-8674. doi: 10.1016/j.cell.2017.11.010.

33. Sebastien Taurin, Reem Alzahrani, Sahar Aloraibi, Layal Ashi, Rawan Alharmi, and Noora Hassani. Patient-derived tumor organoids: A preclinical platform for personalized cancer therapy. Translational Oncology, 51:102226, 2025. ISSN 1936-5233. doi: 10.1016/j.tranon.2024.102226.

34. Yuan-Hung Lo, Kasper Karlsson, and Calvin J. Kuo. Applications of organoids for cancer biology and precision medicine. Nature Cancer, 1(8):761–773, Aug 2020. ISSN 2662-1347. doi: 10.1038/s43018-020-0102-y.

35. Charlotte Bouquerel, Anastasiia Dubrova, Isabella Hofer, Duc T. T. Phan, Moencopi Bernheim, Ségolène Ladaigue, Charles Cavaniol, Danilo Maddalo, Luc Cabel, Fatima Mechta-Grigoriou, Claire Wilhelm, Gérard Zalcman, Maria Carla Parrini, and Stéphanie Descroix. Bridging the gap between tumor-on-chip and clinics: a systematic review of 15 years of studies. Lab Chip, 23: 3906–3935, 2023. doi: 10.1039/D3LC00531C.

36. Hsieh-Fu Tsai, Alen Trubelja, Amy Q. Shen, and Gang Bao. Tumour-on-a-chip: microfluidic models of tumour morphology, growth and microenvironment. J. R. Soc. Interface, 14, 2017.

37. Annika Johnson, Samuel Reimer, Ryan Childres, Grace Cupp, Tia C. L. Kohs, Owen J. T. McCarty, and Youngbok (Abraham) Kang. The applications and challenges of the development of in vitro tumor microenvironment chips. Cellular and Molecular Bioengineering, 16 (1):3–21, Feb 2023. ISSN 1865-5033. doi: 10.1007/s12195-022-00755-7.

38. Marie Nguyen, Adele De Ninno, Arianna Mencattini, Fanny Mermet-Meillon, Giulia Fornabaio, Sophia S. Evans, Mélissande Cossutta, Yasmine Khira, Weijing Han, Philémon Sirven, Floriane Pelon, Davide Di Giuseppe, Francesca Romana Bertani, Annamaria Gerardino, Ayako Yamada, Stéphanie Descroix, Vassili Soumelis, Fatima Mechta-Grigoriou, Gérard Zalcman, Jacques Camonis, Eugenio Martinelli, Luca Businaro, and Maria Carla Parrini. Dissecting effects of anti-cancer drugs and cancer-associated fibroblasts by onchip reconstitution of immunocompetent tumor microenvironments. Cell Reports, 25(13): 3884–3893.e3, Dec 2018. ISSN 2211-1247. doi: 10.1016/j.celrep.2018.12.015.

39. Irina Veith, Martin Nurmik, Arianna Mencattini, Isabelle Damei, Christine Lansche, Solenn Brosseau, Giacomo Gropplero, Stéphanie Corgnac, Joanna Filippi, Nicolas Poté, Edouard Guenzi, Anaïs Chassac, Pierre Mordant, Jimena Tosello, Christine Sedlik, Eliane Piaggio, Nicolas Girard, Jacques Camonis, Hamasseh Shirvani, Fathia Mami-Chouaib, Fatima Mechta-Grigoriou, Stéphanie Descroix, Eugenio Martinelli, Gérard Zalcman, and Maria Carla Parrini. Assessing personalized responses to anti-pd-1 treatment using patientderived lung tumor-on-chip. Cell Reports Medicine, 5(5): 101549, 2024. ISSN 2666-3791. doi: 10.1016/j.xcrm.2024.101549.

40. Martin Nurmik, Ségolène Ladaigue, Isabella Hofer, Auriane Debache, Irina Veith, Fatima Mechta-Grigoriou, Stephanie Descroix, Gerard Zalcman, and Maria Carla Parrini. Protocol for the biofabrication of immunocompetent tumor-on-chip models from patient solid tumors for assessment of anticancer therapies. STAR Protocols, 6(3): 103895, 2025. ISSN 2666-1667. doi: 10.1016/j.xpro.2025.103895.

41. Lisa F. Horowitz, Adan D. Rodriguez, Allan Au-Yeung, Kevin W. Bishop, Lindsey A. Barner, Gargi Mishra, Aashik Raman, Priscilla Delgado, Jonathan T. C. Liu, Taranjit S. Gujral, Mehdi Mehrabi, Mengsu Yang, Robert H. Pierce, and Albert Folch. Microdissected “cuboids” for microfluidic drug testing of intact tissues. Lab Chip, 21: 122–142, 2021. doi: 10.1039/D0LC00801J.

42. Sanjiban Chakrabarty, William F. Quiros-Solano, Maayke M.P. Kuijten, Ben Haspels, Sandeep Mallya, Calvin Shun Yu Lo, Amr Othman, Cinzia Silvestri, Anja van de Stolpe, Nikolas Gaio, Hanny Odijk, Marieke van de Ven, Corrina M.A. de Ridder, Wytske M. van Weerden, Jos Jonkers, Ronald Dekker, Nitika Taneja, Roland Kanaar, and Dik C. van Gent. A microfluidic cancer-on-chip platform predicts drug response using organotypic tumor slice culture. Cancer Research, 82(3):510–520, Feb 2022. ISSN 0008-5472. doi: 10.1158/0008-5472.CAN-21-0799.

43. Marjolijn M. Ladan, Titia G. Meijer, Nicole S. Verkaik, Cecile de Monye, Linetta B. Koppert, Esther Oomen-de Hoop, Carolien H. M. van Deurzen, Roland Kanaar, Julie Nonnekens, Dik C. van Gent, and Agnes Jager. Proof-of-concept study linking ex vivo sensitivity testing to neoadjuvant anthracycline-based chemotherapy response in breast cancer patients. npj Breast Cancer, 9(1):80, Sep 2023. ISSN 2374-4677. doi: 10.1038/s41523-023-00583-6.

44. Louis Jun Ye Ong, Shumei Chia, Stephen Qi Rong Wong, Xiaoqian Zhang, Huiwen Chua, Jia Min Loo, Wei Yong Chua, Clarinda Chua, Emile Tan, Hannes Hentze, Iain Beehuat Tan, Ramanuj DasGupta, and Yi-Chin Toh. A comparative study of tumour-on-chip models with patient-derived xenografts for predicting chemotherapy efficacy in colorectal cancer patients. Frontiers in Bioengineering and Biotechnology, Volume 10 - 2022, 2022. ISSN 2296-4185. doi: 10.3389/fbioe.2022.952726.

45. Eliana Steinberg, Roy Friedman, Yoel Goldstein, Nethanel Friedman, Ofer Beharier, Jonathan Abraham Demma, Gideon Zamir, Ayala Hubert, and Ofra Benny. A fully 3d-printed versatile tumor-on-a-chip allows multi-drug screening and correlation with clinical outcomes for personalized medicine. Communications Biology, 6(1):1157, Nov 2023. ISSN 2399-3642. doi: 10.1038/s42003-023-05531-5.

46. Meiling Zhang and Bin Zhang. Extracellular matrix stiffness: mechanisms in tumor progression and therapeutic potential in cancer. Experimental Hematology & Oncology, 14(1):54, Apr 2025. ISSN 2162-3619. doi: 10.1186/s40164-025-00647-2.

47. Jason J. Northey, Alexander S. Barrett, Irene Acerbi, Mary-Kate Hayward, Stephanie Talamantes, Ivory S. Dean, Janna K. Mouw, Suzanne M. Ponik, Jonathon N. Lakins, Po-Jui Huang, Junmin Wu, Quanming Shi, Susan Samson, Patricia J. Keely, Rita A. Mukhtar, Jan T. Liphardt, John A. Shepherd, E. Shelley Hwang, Yunn-Yi Chen, Kirk C. Hansen, Laurie E. Littlepage, and Valerie M. Weaver. Stiff stroma increases breast cancer risk by inducing the oncogene znf217. The Journal of Clinical Investigation, 130(11):5721–5737, 11 2020. doi: 10.1172/JCI129249.

48. Su-Yeong Jeong, Ji-Hyun Lee, Yoojin Shin, Seok Chung, and Hyo-Jeong Kuh. Co-culture of tumor spheroids and fibroblasts in a collagen matrix-incorporated microfluidic chip mimics reciprocal activation in solid tumor microenvironment. PLOS ONE, 11(7):1–17, 07 2016. doi: 10.1371/journal.pone.0159013.

49. Pilar Alamán-Díez, Carlos Borau, Pedro Enrique Guerrero, Hippolyte Amaveda, Mario Mora, José María Fraile, Elena García-Gareta, José Manuel García-Aznar, and María Ángeles Pérez. Collagen-laponite nanoclay hydrogels for tumor spheroid growth. Biomacromolecules, 24(6):2879–2891, Jun 2023. ISSN 1525-7797. doi: 10.1021/acs.biomac.3c00257.

50. Michela Anna Polidoro, Erika Ferrari, Cristiana Soldani, Barbara Franceschini, Giuseppe Saladino, Arianna Rosina, Andrea Mainardi, Francesca D’Autilia, Nicola Pugliese, Guido Costa, Matteo Donadon, Guido Torzilli, Simona Marzorati, Marco Rasponi, and Ana Lleo. Cholangiocarcinoma-on-a-chip: A human 3d platform for personalised medicine. JHEP Reports, 6(1): 100910, 2024. ISSN 2589-5559. doi: 10.1016/j.jhepr.2023.100910.

51. Soraya Hernández-Hatibi, Pedro Enrique Guerrero, José Manuel García-Aznar, and Elena García-Gareta. Polydopamine interfacial coating for stable tumor-on-a-chip models: Application for pancreatic ductal adenocarcinoma. Biomacromolecules, 25(8):5169–5180, Aug 2024. ISSN 1525-7797. doi: 10.1021/acs.biomac.4c00551.

52. Funda Meric-Bernstam, Michael W. Lloyd, Soner Koc, Yvonne A. Evrard, Lisa M. McShane, Michael T. Lewis, Kurt W. Evans, Dali Li, Lawrence Rubinstein, Alana Welm, II Dean, Dennis A., Anuj Srivastava, Jeffrey W. Grover, Min J. Ha, Huiqin Chen, Xuelin Huang, Kaushik Varadarajan, Jing Wang, Jack A. Roth, Bryan Welm, Ramaswamy Govinden, Li Ding, Salma Kaochar, Nicholas Mitsiades, Luis Carvajal-Carmona, Meenhard Herylyn, Michael A. Davies, Geoffrey I. Shapiro, Ryan Fields, Jose G. Trevino, Joshua C. Harrell, NCI PDXNet Consortium, James H. Doroshow, Jeffrey H. Chuang, and Jeffrey A. Moscow. Assessment of patient-derived xenograft growth and antitumor activity: The nci pdxnet consensus recommendations. Molecular Cancer Therapeutics, 23(7):924–938, Jul 2024. ISSN 1535-7163. doi: 10.1158/1535-7163.MCT-23-0471.

53. Dario Trapani, Diogo Martins-Branco, Giuseppe Curigliano, Alessandra Gennari, George Pentheroudakis, and Nadia Harbeck. Updated treatment recommendations for systemic treatment: from the esmo metastatic breast cancer living guideline^†^. Annals of Oncology, october 2025. ISSN 0923-7534. doi: 10.1016/j.annonc.2025.07.017.

54. Dane R. Liston and Myrtle Davis. Clinically relevant concentrations of anticancer drugs: A guide for nonclinical studies. Clinical Cancer Research, 23(14):3489–3498, Jul 2017. ISSN 1078-0432. doi: 10.1158/1078-0432.CCR-16-3083.

55. Mark C. Pettinato. Introduction to antibody-drug conjugates. Antibodies, 10(4), 2021. ISSN 2073-4468. doi: 10.3390/antib10040042.

56. Shunji Takahashi, Masato Karayama, Masato Takahashi, Junichiro Watanabe, Hironobu Minami, Noboru Yamamoto, Ichiro Kinoshita, Chia-Chi Lin, Young-Hyuck Im, Issei Achiwa, Emi Kamiyama, Yasuyuki Okuda, Caleb Lee, and Yung-Jue Bang. Pharmacokinetics, safety, and efficacy of trastuzumab deruxtecan with concomitant ritonavir or itraconazole in patients with her2-expressing advanced solid tumors. Clinical Cancer Research, 27(21):5771–5780, Nov 2021. ISSN 1078-0432. doi: 10.1158/1078-0432.CCR-21-1560.

57. Ophelia Yin, Yuan Xiong, Seiko Endo, Kazutaka Yoshihara, Tushar Garimella, Malaz Abu-Tarif, Russ Wada, and Frank LaCreta. Population pharmacokinetics of trastuzumab deruxtecan in patients with her2-positive breast cancer and other solid tumors. Clinical Pharmacology & Therapeutics, 109(5): 1314–1325, 2021. doi: 10.1002/cpt.2096.

58. Christina Vasalou, Theresa A. Proia, Laura Kazlauskas, Anna Przybyla, Matthew Sung, Srinivas Mamidi, Kim Maratea, Matthew Griffin, Rebecca Sargeant, Jelena Urosevic, Anton I. Rosenbaum, Jiaqi Yuan, Krishna C. Aluri, Diane Ramsden, Niresh Hariparsad, Rhys D.O. Jones, and Jerome T. Mettetal. Quantitative evaluation of trastuzumab deruxtecan pharmacokinetics and pharmacodynamics in mouse models of varying degrees of her2 expression. CPT: Pharmacometrics & Systems Pharmacology, 13(6): 994–1005, 2024. doi: 10.1002/psp4.13133.

59. Peter Schmid, Javier Cortes, Lajos Pusztai, Heather McArthur, Sherko Kümmel, Jonas Bergh, Carsten Denkert, Yeon Hee Park, Rina Hui, Nadia Harbeck, Masato Takahashi, Theodoros Foukakis, Peter A. Fasching, Fatima Cardoso, Michael Untch, Liyi Jia, Vassiliki Karantza, Jing Zhao, Gursel Aktan, Rebecca Dent, and Joyce O’Shaughnessy. Pembrolizumab for early triple-negative breast cancer. New England Journal of Medicine, 382 (9):810–821, 2020. doi: 10.1056/NEJMoa1910549.

60. Ergang Wang, Kun Xiang, Yun Zhang, and Xiao-Fan Wang. Patient-derived organoids (pdos) and pdo-derived xenografts (pdoxs): New opportunities in establishing faithful preclinical cancer models. Journal of the National Cancer Center, 2(4): 263–276, 2022. ISSN 2667-0054. doi: 10.1016/j.jncc.2022.10.001.

61. Miguel Martín, Atanasio Pandiella, Emilio Vargas-Castrillón, Elena Díaz-Rodríguez, Teresa Iglesias-Hernangómez, Concha Martínez Cano, Inés Fernández-Cuesta, Elena Winkow, and Maria Francesca Perelló. Trastuzumab deruxtecan in breast cancer. Critical Reviews in Oncology/Hematology, 198:104355, 2024. ISSN 1040-8428. doi: 10.1016/j.critrevonc.2024.104355.

62. Li-Chung Tsao, John S. Wang, Xingru Ma, Sirajbir Sodhi, Joey V. Ragusa, Bushangqing Liu, Jason McBane, Tao Wang, Junping Wei, Cong-Xiao Liu, Xiao Yang, Gangjun Lei, Ivan Spasojevic, Ping Fan, Timothy N. Trotter, Michael Morse, Herbert Kim Lyerly, and Zachary C. Hartman. Effective extracellular payload release and immunomodulatory interactions govern the therapeutic effect of trastuzumab deruxtecan (t-dxd). Nature Communications, 16 (1):3167, Apr 2025. ISSN 2041-1723. doi: 10.1038/s41467-025-58266-8.

63. Yusuke Ogitani, Katsunobu Hagihara, Masataka Oitate, Hiroyuki Naito, and Toshinori Agatsuma. Bystander killing effect of ds-8201a, a novel anti-human epidermal growth factor receptor 2 antibody–drug conjugate, in tumors with human epidermal growth factor receptor 2 heterogeneity. Cancer Science, 107(7): 1039–1046, 2016. doi: 10.1111/cas.12966.

64. Elisabetta Marangoni, Cécile Laurent, Florence Coussy, Rania El-Botty, Sophie Château-Joubert, Jean-Luc Servely, Ludmilla de Plater, Franck Assayag, Ahmed Dahmani, Elodie Montaudon, Fariba Nemati, Justine Fleury, Sophie Vacher, David Gentien, Audrey Rapinat, Pierre Foidart, Nor Eddine Sounni, Agnès Noel, Anne Vincent-Salomon, Marick Lae, Didier Decaudin, Sergio Roman-Roman, Ivan Bièche, Martine Piccart, and Fabien Reyal. Capecitabine efficacy is correlated with tymp and rb1 expression in pdx established from triple-negative breast cancers. Clinical Cancer Research, 24(11):2605–2615, May 2018. ISSN 1078-0432. doi: 10.1158/1078-0432.CCR-17-3490.

65. Petra ter Brugge, Sarah C. Moser, Ivan Bièche, Petra Kristel, Sabrina Ibadioune, Alexandre Eeckhoutte, Roebi de Bruijn, Eline van der Burg, Catrin Lutz, Stefano Annunziato, Julian de Ruiter, Julien Masliah Planchon, Sophie Vacher, Laura Courtois, Rania El-Botty, Ahmed Dahmani, Elodie Montaudon, Ludivine Morisset, Laura Sourd, Léa Huguet, Heloise Derrien, Fariba Nemati, Sophie Chateau-Joubert, Thibaut Larcher, Anne Salomon, Didier Decaudin, Fabien Reyal, Florence Coussy, Tatiana Popova, Jelle Wesseling, Marc-Henri Stern, Jos Jonkers, and Elisabetta Marangoni. Homologous recombination deficiency derived from whole-genome sequencing predicts platinum response in triplenegative breast cancers. Nature Communications, 14(1):1958, Apr 2023. ISSN 2041-1723. doi: 10.1038/s41467-023-37537-2.

66. Florence Coussy, Rania El-Botty, Sophie Château-Joubert, Ahmed Dahmani, Elodie Montaudon, Sophie Leboucher, Ludivine Morisset, Pierre Painsec, Laura Sourd, Léa Huguet, Fariba Nemati, Jean-Luc Servely, Thibaut Larcher, Sophie Vacher, Adrien Briaux, Cécile Reyes, Philippe La Rosa, Georges Lucotte, Tatiana Popova, Pierre Foidart, Nor Eddine Sounni, Agnès Noel, Didier Decaudin, Laetitia Fuhrmann, Anne Salomon, Fabien Reyal, Christopher Mueller, Petra Ter Brugge, Jos Jonkers, Marie-France Poupon, Marc-Henri Stern, Ivan Bièche, Yves Pommier, and Elisabetta Marangoni. Brcaness, slfn11, and rb1 loss predict response to topoisomerase i inhibitors in triple-negative breast cancers. Science Translational Medicine, 12(531):eaax2625, 2020. doi: 10.1126/scitranslmed.aax2625.

67. F. Coussy, R. El Botty, M. Lavigne, C. Gu, L. Fuhrmann, A. Briaux, L. de Koning, A. Dahmani, E. Montaudon, L. Morisset, L. Huguet, L. Sourd, P. Painsec, S. Chateau-Joubert, T. Larcher, S. Vacher, S. Melaabi, A. Vincent Salomon, E. Marangoni, and I. Bieche. Combination of pi3k and mek inhibitors yields durable remission in pdx models of pik3ca-mutated metaplastic breast cancers. Journal of Hematology & Oncology, 13(1):13, Feb 2020. ISSN 1756-8722. doi: 10.1186/s13045-020-0846-y.

