## Supplementary material for "Tumor-on-Chip as a Personalised Platform for Rapid Drug-Testing in Breast Cancer": Supp. Info.

### Supplementary Figures

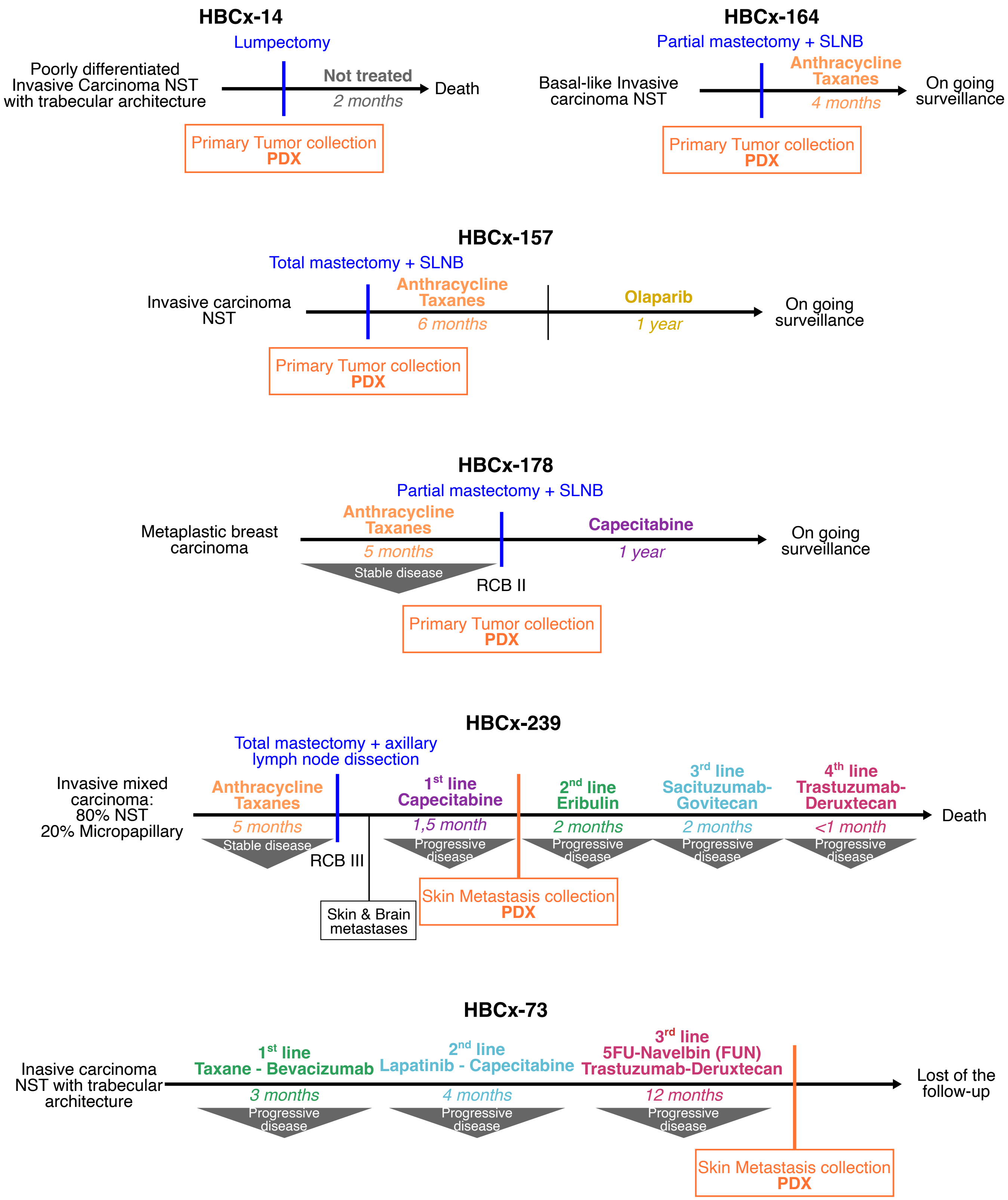


**Fig. Supp. 1. Characteristics of breast cancer models.** Overview of the key features and clinical treatment history of patients from whom PDX models were derived. For each treatment, the corresponding progression-free survival (PFS) and best response according to RECIST v1.1 criteria are indicated.
NST = no special type; RCB = residual cancer burden; SLNB = sentinel lymph node biopsy.

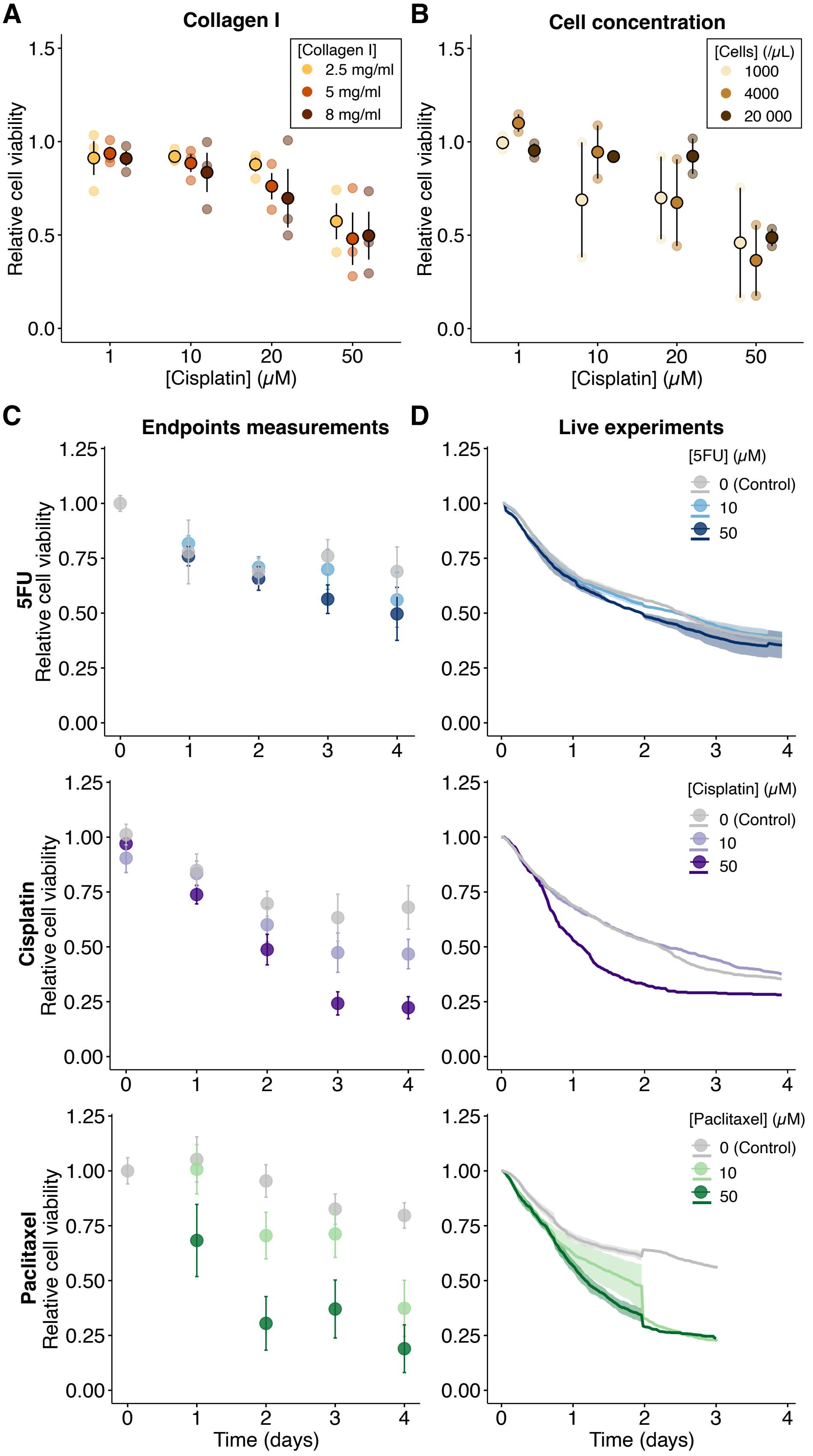


**Fig. Supp. 2. Calibration of the chips' features.** Graphs representing the impact of (A) collagen I concentration and (B) cell concentration on cell viability after 3 days of exposure to cisplatin 50 µM using the HBCx-14 model. For (B), a collagen concentration of 2.5 mg/mL was selected. Each data point represents an individual chip; central points denote the mean of 3 chips and the error bar corresponds to the SD (C-D). Kinetics of cell viability measured in different chips for each time point, placed into an incubator (C), and measured longitudinally in one chip under a spinning disk microscope (D). In (C), each point represents the mean of at least 3 chips, and the error bar corresponds to the SD. In (D), each curve represents the mean of 2 chips. The full shade area corresponds to the SD.

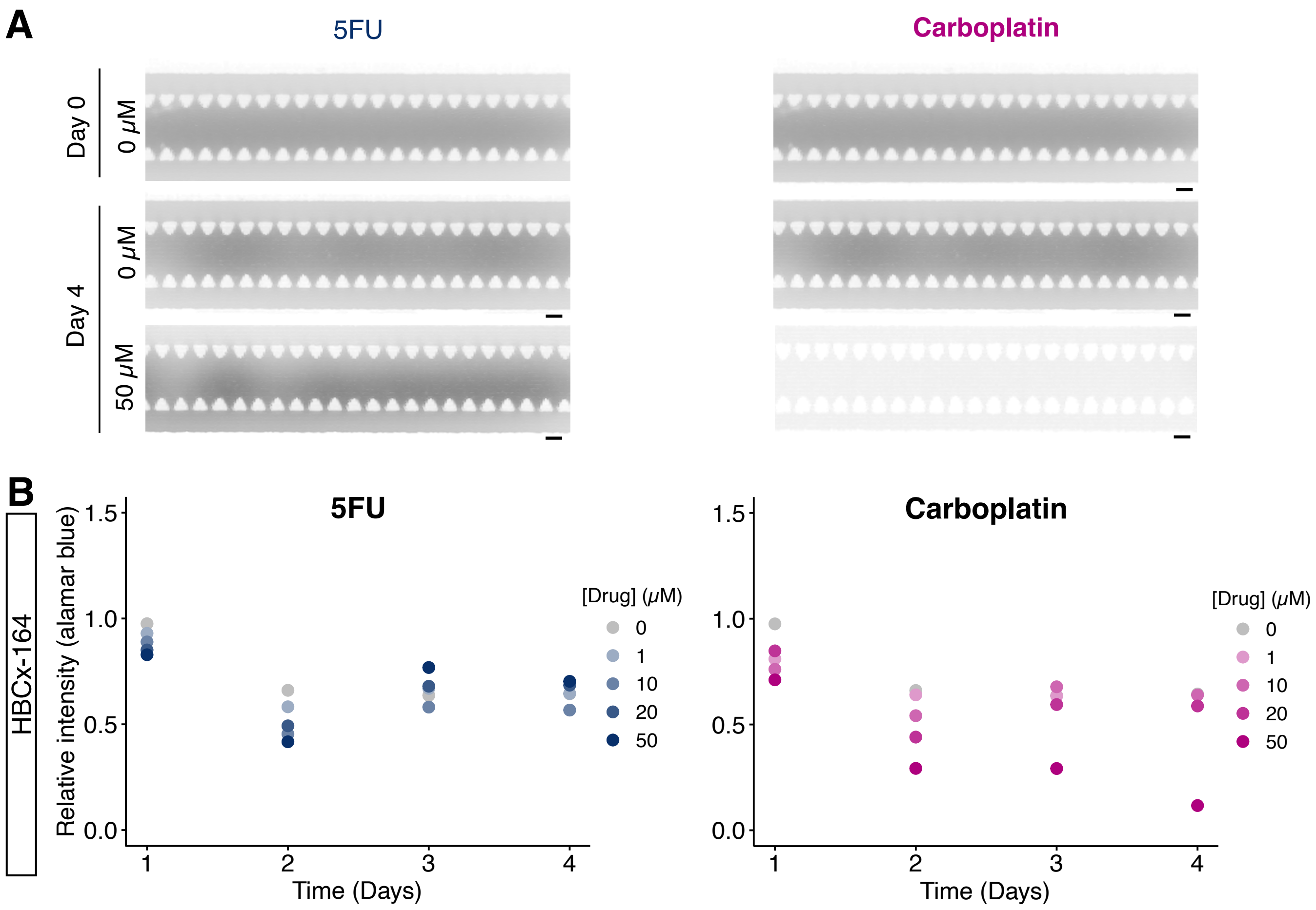


**Fig. Supp. 3. Metabolic assay assessing cytotoxicity** (A) Representative images of the metabolic assay (AlamarBlue dye) performed on the HBCx-14 breast cancer model treated with 5FU (top) and carboplatin (bottom). Scale bar: 100 µm. (B) Graphs representing the cytotoxic effect of 5FU and carboplatin on HBCx-14 tumor cells (N=1).

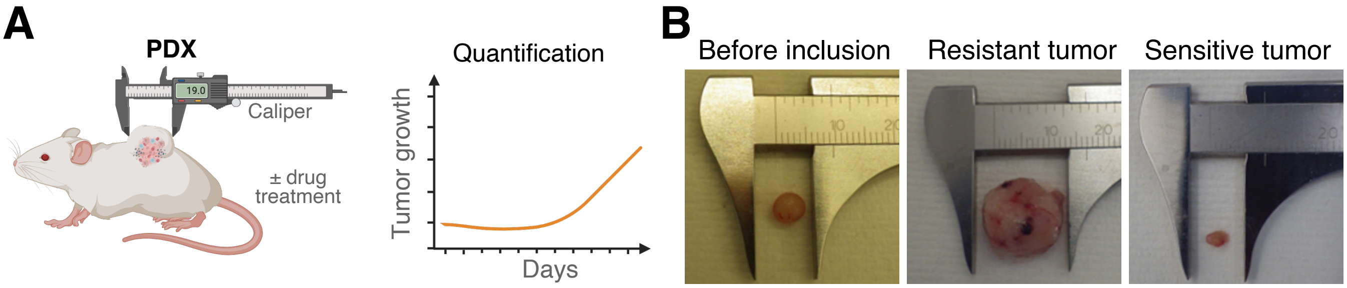


**Fig. Supp. 4. PDX tumor growth.** (A) Schematics of PDX experimental setup. Tumor growth was measured twice a week using a caliper. B) Representative images of tumors before engraftement (left), and after excision following treatment exposure showing models classified as resistant (middle) or sensitive (right) to treatment.

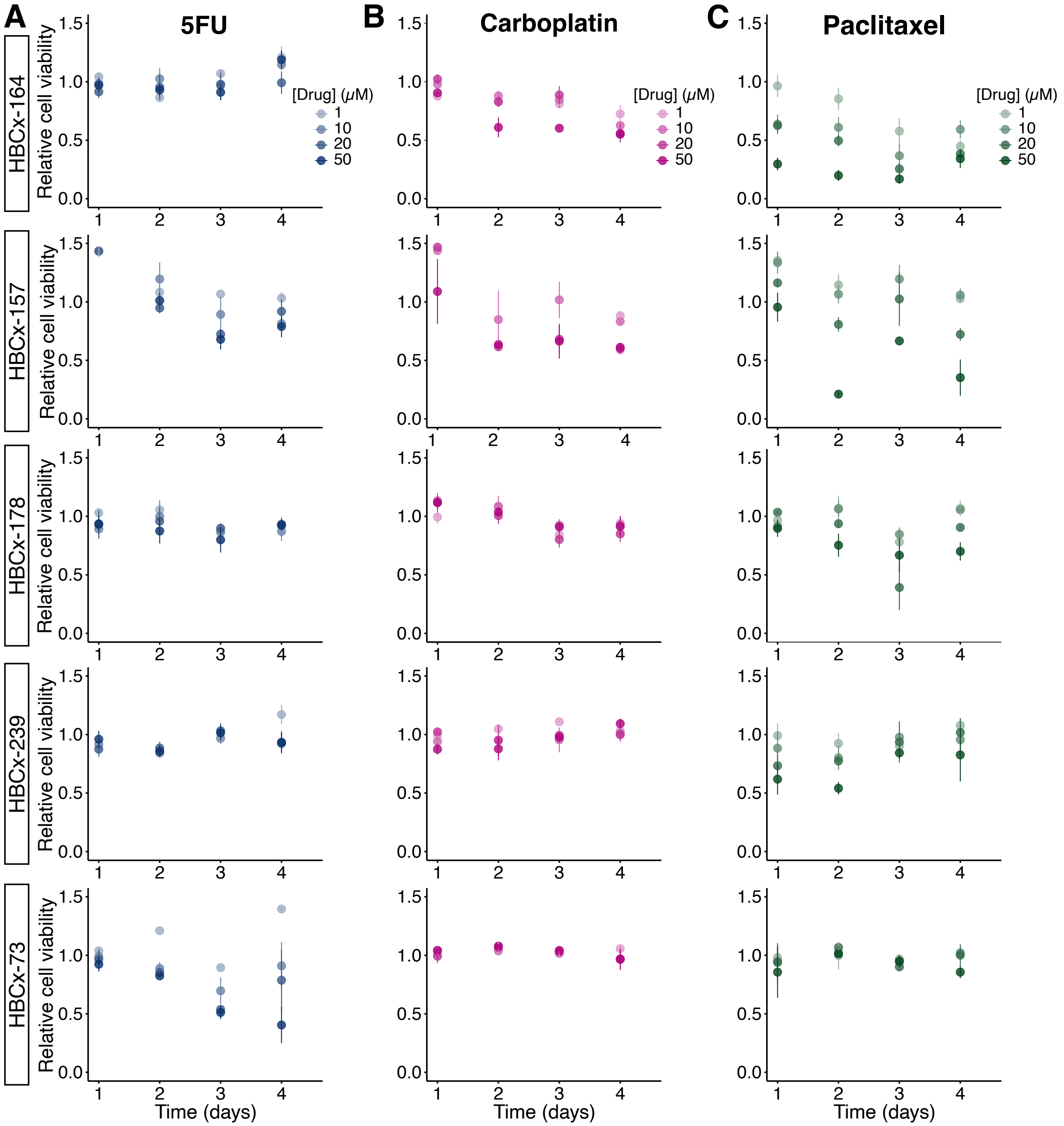


**Fig. Supp. 5. Time-course comparison of responses to multiple drugs across different models.** Graphs representing the kinetics of cell viability over time. Each model was treated with (A) 5FU, (B) carboplatin, and (C) paclitaxel from 1 to 50 µM. In all conditions, individual data was normalized by the initial point (Day 0), and with the control condition (concentration = 0) at each day. Each point represents the mean of at least 2 independent experiments, and the error bar corresponds to a SD. All statistics were performed using the Wilcoxon test. ns = p-value > 0.05, * = p-value ≤ 0.05, ** = p-value ≤ 0.01, *** = p-value ≤ 0.001, **** = p-value ≤ 0.0001.

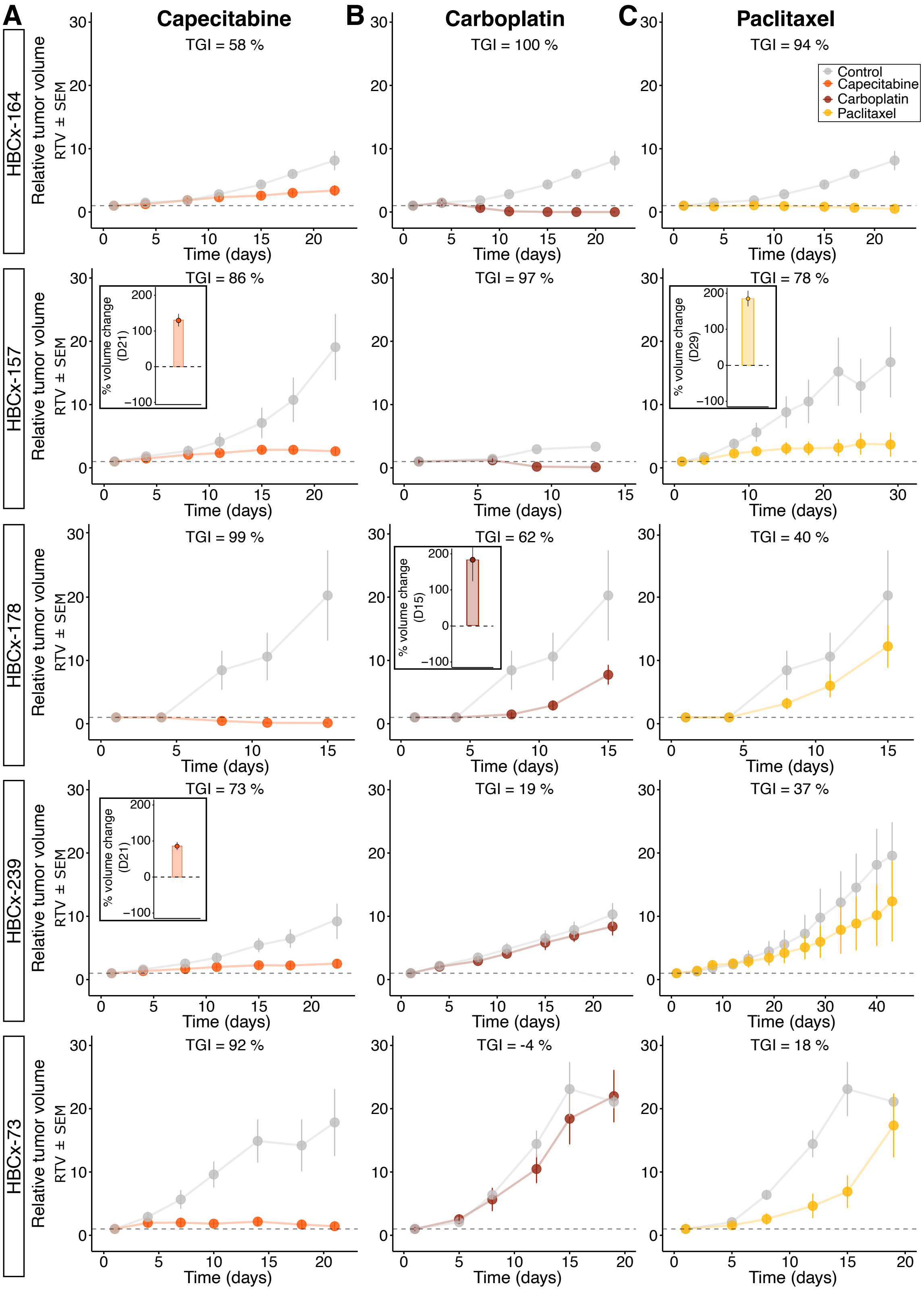


**Fig. Supp. 6. Kinetics of PDX tumor growth *in vivo*.** PDX models were treated with the indicated agents. Tumor volumes (mm³) were measured longitudinally; each point represents the mean of at least three independent mice, and error bars indicate the SEM. Treated groups are shown as colored curves and untreated controls as a gray curve. Treatment regimens were: (A) capecitabine, 540 mg/kg, five times per week per os; (B) carboplatin, 90 mg/kg, once every three weeks IP; and (C) paclitaxel, 25 mg/kg, once per week IP. For intermediate tumor growth inhibition (TGI, 60–90%), histograms show the relative mean tumor volume change (%VC) of the treated group, illustrating resistance to treatment.

| Tumor | Model | Metric | 5FU / Capecitabine | Carboplatin | Paclitaxel | Cisplatin |
|---|---|---|---|---|---|---|
| **HBCx-14** | PDX | TGI | 41% (D43) | 100% (D22) | 92% (D22) | 99% (D22) |
| | | % volume change | 584% (D43) | -100% (D22) | -100% (D22) | -100% (D22) |
| | ToC | Relative cell viability | 0.72 (D4) | 0.51 (D4) | 0.33 (D4) | 0.24 (D4) |
| **HBCx-164** | PDX | TGI | 58% (D22) | 100% (D22) | 94% (D22) | |
| | | % volume change | 284% (D22) | -100% (D22) | -64% (D22) | |
| | ToC | Relative cell viability | 1.19 (D4) | 0.55 (D4) | 0.34 (D4) | |
| **HBCx-157** | PDX | TGI | **86% (D22)** | 97% (D13) | **78% (D29)** | |
| | | % volume change | **217% (D25)** | -93% (D27) | **263% (D29)** | |
| | ToC | Relative cell viability | 0.79 (D4) | 0.61 (D4) | 0.35 (D4) | |
| **HBCx-178** | PDX | TGI | 99% (D15) | **62% (D15)** | 39% (D15) | |
| | | % volume change | -84% (D15) | **682% (D15)** | 1129% (D15) | |
| | ToC | Relative cell viability | 0.93 (D4) | 0.91 (D4) | 0.70 (D4) | |
| **HBCx-239** | PDX | TGI | **73% (D22)** | 19% (D22) | 37% (D43) | |
| | | % volume change | **147% (D22)** | 829% (D22) | 1265% (D43) | |
| | ToC | Relative cell viability | 0.93 (D4) | 1.09 (D4) | 0.83 (D4) | |
| **HBCx-73** | PDX | TGI | 92% (D21) | -4% (D19) | 18% (D19) | |
| | | % volume change | -25% (D28) | 1878% (D19) | 1444% (D19) | |
| | ToC | Relative cell viability | 0.40 (D4) | 0.97 (D4) | 0.86 (D4) | |

% Volume change:
Progression disease (PD): > 35%
Stable disease (SD): -50% to 35%
Partial response (PR): -94% to -51%
Complete response (CR): -95% to -100%

Resistant: TGI < 60%
Intermediate: 60% < TGI < 90%
Sensitive: TGI > 90%

PD and TGI < 90%
SD, PR or CR and TGI > 90%

**Fig. Supp. 7. Summary table of the PDX and ToC.** Features of PDX and ToC.

**A**

| Models | HBCx-14 #1 | HBCx-14 #2 | HBCx-14 #3 | HBCx-178 #1 | HBCx-178 #2 | HBCx-178 #3 | HBCx-239 #1 | HBCx-239 #2 | HBCx-239 #3 | HBCx-73 #1 |
|---|---|---|---|---|---|---|---|---|---|---|
| Dimensions of the biopsy | 12x1.02 mm | 11x1.02 mm | 8x1.02 mm | 20x1.02 mm | 20x1.02 mm | 10x1.02 mm | 10x1.02 mm | 20x1.02 mm | 20x1.02 mm | 20x1.02 mm |
| # Biopsies | 4 | 4 | 4 | 3 | 3 | 4 | 4 | 3 | 3 | 3 |
| # Total of Cells | 300 000 | 250 000 | 125 000 | 720 000 | 350 000 | 100 000 | 100 000 | 200 000 | 100 000 | 641 000 |

**B**

| Patient | #1 | #2 | #3 | #4 | #5 | #6 |
|---|---|---|---|---|---|---|
| # Total of Cells | 110 000 | 850 000 | 1 000 000 | 5 760 000 | 2 565 000 | 800 000 |
| Type of the sample | Resection | Resection | Resection | Resection | Resection | Biopsy |

**Fig. Supp. 8. Features of the biopsies.** Characteristics of the biopsies performed on (A) PDX tumors and (B) patient samples.
